# Microbial responses to stress cryptically alter natural selection on plants

**DOI:** 10.1101/2024.02.09.579697

**Authors:** Lana G. Bolin, Jennifer A. Lau

## Abstract

- Microbial communities can rapidly respond to stress, meaning plants may encounter altered soil microbial communities in stressful environments. These altered microbial communities may then affect natural selection on plants. Because stress can cause lasting changes to microbial communities, microbes may also cause legacy effects on plant selection that persist even after the stress ceases.
- To explore how microbial responses to stress and persistent microbial legacy effects of stress affect natural selection, we grew *Chamaecrista fasciculata* plants in stressful (salt, herbicide, or herbivory) or non-stressful conditions with microbes that had experienced each of these environments in the previous generation.
- Microbial community responses to stress generally counteracted the effects of stress itself on plant selection, thereby weakening the strength of stress as a selective agent. Microbial legacy effects of stress altered plant selection in non-stressful environments, suggesting that stress-induced changes to microbes may continue to affect selection after stress is lifted.
- These results suggest that soil microbes may play a cryptic role in plant adaptation to stress, potentially reducing the strength of stress as a selective agent and altering the evolutionary trajectory of plant populations.

## INTRODUCTION

Environmental stress commonly alters patterns of natural selection in plants (e.g., Stanton *et al*., 2000), and it is often assumed that the stress itself is driving this change in selection. However, soil microbial communities can also affect natural selection on plants (Lau & Lennon, 2011; Wagner *et al*., 2014; Chaney & Baucom, 2020), and stress may alter soil microbial communities through rapid shifts in community composition or the evolution of key taxa (Elena & Lenski, 2003; Graves *et al*., 2015). Microbes may therefore play a cryptic role in plant evolutionary responses to stress if microbial communities respond to environmental stress, and these new microbial communities then alter the strength and/or direction of natural selection acting on plant traits.

Microbial community responses to stress may affect selection in several ways (Fig. 1). In some cases, the microbial community might not respond strongly to stress, or even if it does, the altered microbial community may not change selection on plant hosts. In these cases, stress itself primarily alters selection as is commonly assumed (Fig. **1a**). However, in other cases shifts in microbial communities may strongly alter selection, and the effects of stress-induced shifts in community composition could equal or even exceed the effects of stress itself. In these cases, ‘stress-adapted’ microbial communities (here used to include both shifts in microbial community composition and microbial evolution in response to stress) are the primary drivers of changes in selection under stress such that shifts in selection are only detected with the stress-adapted microbial community (i.e., microbes may be cryptic drivers of selection on plant traits; Fig. **1b**). In other cases, microbial community responses to stress could counteract the effects of stress, thereby weakening the overall effect of stress on selection (Fig. **1c**). In addition to influencing plant selection under contemporary stress, stress-adapted microbes may also influence natural selection for the next generation of plants via a microbial legacy of stress, even if the new generation does not experience stress (Fig. **1d**). Such microbial legacy effects of stress might occur when microbes respond to stress in ways that alter plant selection and these effects persist even after the stress ceases.

**Figure 1.**
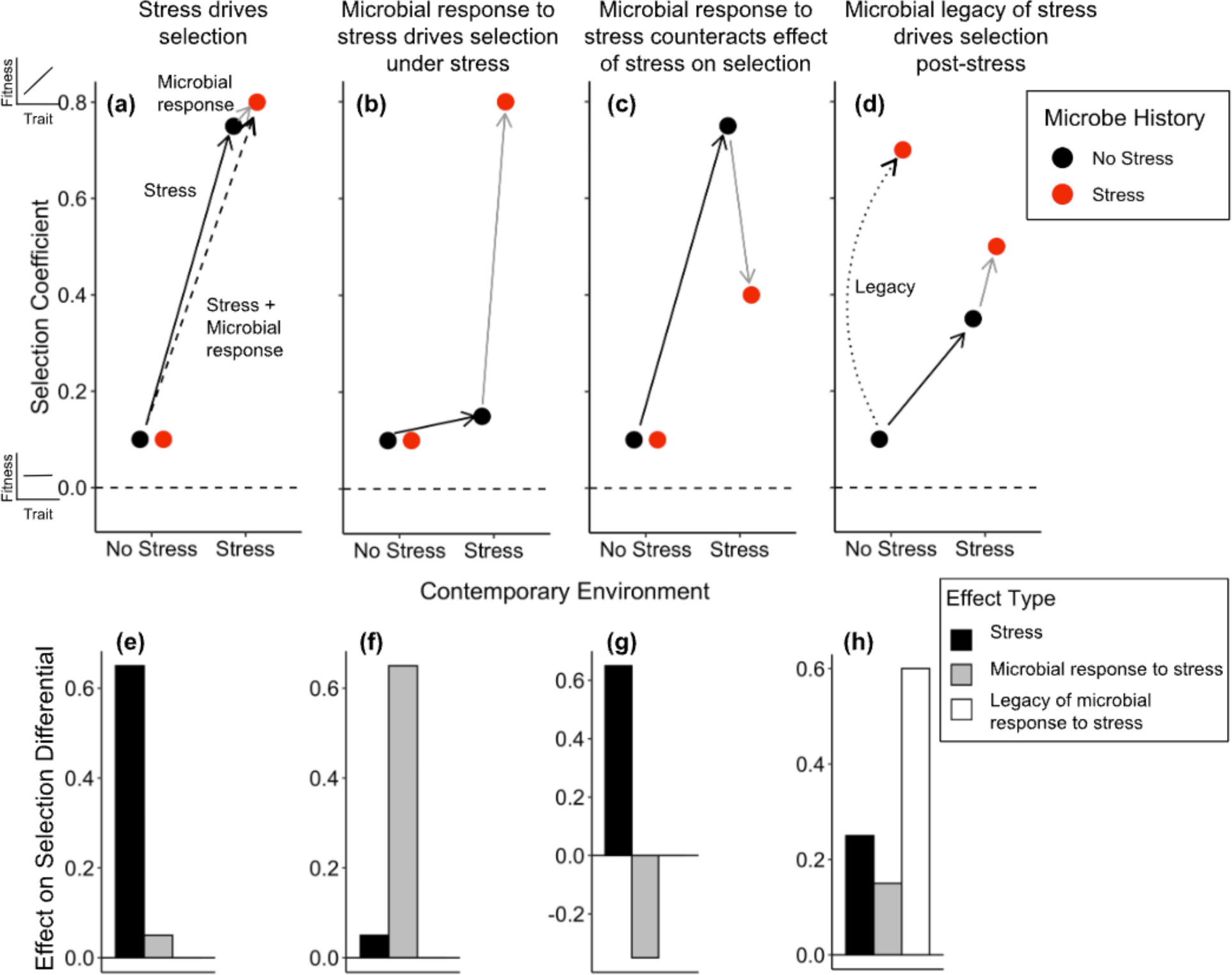
Conceptual figure showing how soil microbial responses to stress (‘Microbe History’) may influence patterns of natural selection (i.e., the selection coefficient) acting on plants in stressful environments. A positive selection coefficient means higher trait values are favored by natural selection, with more positive coefficients indicating stronger positive selection. **(a)** Stress itself may directly alter natural selection acting on plants (i.e., microbial response to stress {‘microbe history’} does not affect selection), or **(b)** stress may alter natural selection on plants by causing rapid changes in the soil microbial community, and these new ‘stress’ microbes then drive changes in plant natural selection observed under stress. **(c)** Alternatively, ‘stress’ microbes could counteract the effect of stress on plant natural selection. **(d)** Finally, a microbial legacy of stress could drive plant natural selection post-stress if ‘stress’ microbial communities remain intact in the soil and alter selection for the next generation of plants who never experienced the stress. The effect of stress itself is shown as a solid arrow, the combined effect of stress and microbial community response to stress is shown as a dashed arrow, the effect of microbial responses to stress is shown as a gray arrow, and the microbial legacy effect of stress is shown as a curved dotted arrow. **(e-h)** The effect of stress on selection (black bars), the effect of microbial community responses to stress on selection (gray bars), and the legacy effect of microbial responses to stress on selection (white bar) are shown for each hypothetical case as calculated using Equations 1-3 in the main text (these bar graphs are analogous to those shown in Figure 4). In this example selection tends to be stronger under stress than in the absence of stress, but stress could also weaken or change the direction of natural selection. Note: the effect of stress itself may also include extremely rapid changes in microbial community composition.

These legacy effects may lead to longer-lasting effects of stress on patterns of natural selection in plants. In addition to altering the strength or direction of natural selection acting on particular plant traits, soil microbial community responses to stress also might mediate plant evolutionary responses to stress by affecting the opportunity for selection (*I*), that is, the variance in relative fitness (Arnold & Wade, 1984; Crow, 1958; Caruso *et al*., 2017). The opportunity for selection limits the maximum strength of selection that can act on any phenotype in a population (Crow, 1958). Because *I* can be expressed as the ratio of variance in absolute fitness to squared mean absolute fitness (Wade & Shuster, 2005), *I* is predicted to be larger in environments that reduce mean fitness in absolute terms (Arnold & Wade, 1984; Rundle & Vamosi, 1996; Fugère & Hendry, 2018). This is because a better-than-average individual will have higher relative fitness in such environments and a worse-than-average individual will have lower relative fitness, resulting in an increase in variance in relative fitness (Fugère & Hendry, 2018). Therefore, microbial responses to stress may increase *I* if they exacerbate the negative fitness effects of stress for plants (e.g., pathogens dominate in stressful environments), or they may reduce *I* if they buffer plants from stress (e.g., mutualists dominate in stressful environments, or stressful environments favor microbial communities that buffer plants from stress) (e.g., Lau & Lennon, 2012; Fitzpatrick *et al*., 2018; Giauque *et al*., 2019; Allsup & Lankau, 2019; Bolin *et al*., 2022). Microbial community responses to stress also may affect *I* by altering the variance in absolute fitness under stress, which might occur if plant genotypes differ in the benefits (or harm) they receive from the microbial taxa that become more abundant under stress (e.g., Bever *et al*., 1996; Eck *et al*., 2019; Bolin & Lau, 2022).

Here, we investigated the evolutionary consequences of plant interactions with diverse soil microbes under stress. We independently manipulated three stress environments (salt, herbicide, and herbivory stress) and the soil microbes those stress environments select for to test how microbial responses to stress influence plant evolutionary processes. First, we quantified the effects of stress on plant natural selection, how stress-induced changes to the soil microbial community affect plant natural selection, and the legacy effects that result from microbial responses to past stress altering selection for future plant generations growing in non-stressful environments. We compared the strength and direction of these effects across stress environments and plant traits, and we also estimated selection acting through both plant survival and fecundity to examine how microbial effects on selection may differ across plant life history stages. Second, we tested how stress and microbial responses to stress affect *I*, and whether these changes were driven by changes in mean absolute fitness or variance in absolute fitness. Our findings illustrate that, in addition to the well-studied effects of soil microbes on plant ecology (e.g., abundance, community composition, succession; Kardol *et al*. 2006; Kulmatiski *et al*., 2008; van der Putten *et al*., 2013), microbial community responses to stress also may alter plant evolution. As a result, evolutionary effects of stress on plants may be driven as much by microbial community responses to the stress as by the stress itself.

## MATERIALS AND METHODS

### Overview

We tested how stress and microbial responses to stress affect natural selection on plants using the annual legume *Chamaecrista fasciculata* (*Fabaceae; Chamaecrista* hereafter). We collected soil inoculum from experimental plots grown with *Chamaecrista* that had been treated with one of four stress treatments: salt, herbicide, simulated herbivory, or an unstressed control (‘microbe history’ hereafter; n = 3 randomized field plots per stress treatment; Fig. 2). We then planted *Chamaecrista* into mesocosms inoculated with each of these microbial communities in the glasshouse and applied the same four stress treatments (‘contemporary environment’ hereafter) in a full factorial design. We planted an individual from each of 50 full-sib families into each field plot × contemporary environment treatment, resulting in 3 individuals per family per microbe history × contemporary environment treatment (N = 2,400 plants = 3 individuals per family per treatment combination × 50 families × 16 treatment combinations); this family structure allowed us to conduct genotypic selection analyses (see ‘Statistical Analyses’ methods). These 50 plants were spread across two mesocosms with 25 plants per mesocosm due to pot size constraints, and families were randomly assigned to a location within one of these two mesocosms (Fig. 2).

**Figure 2.**
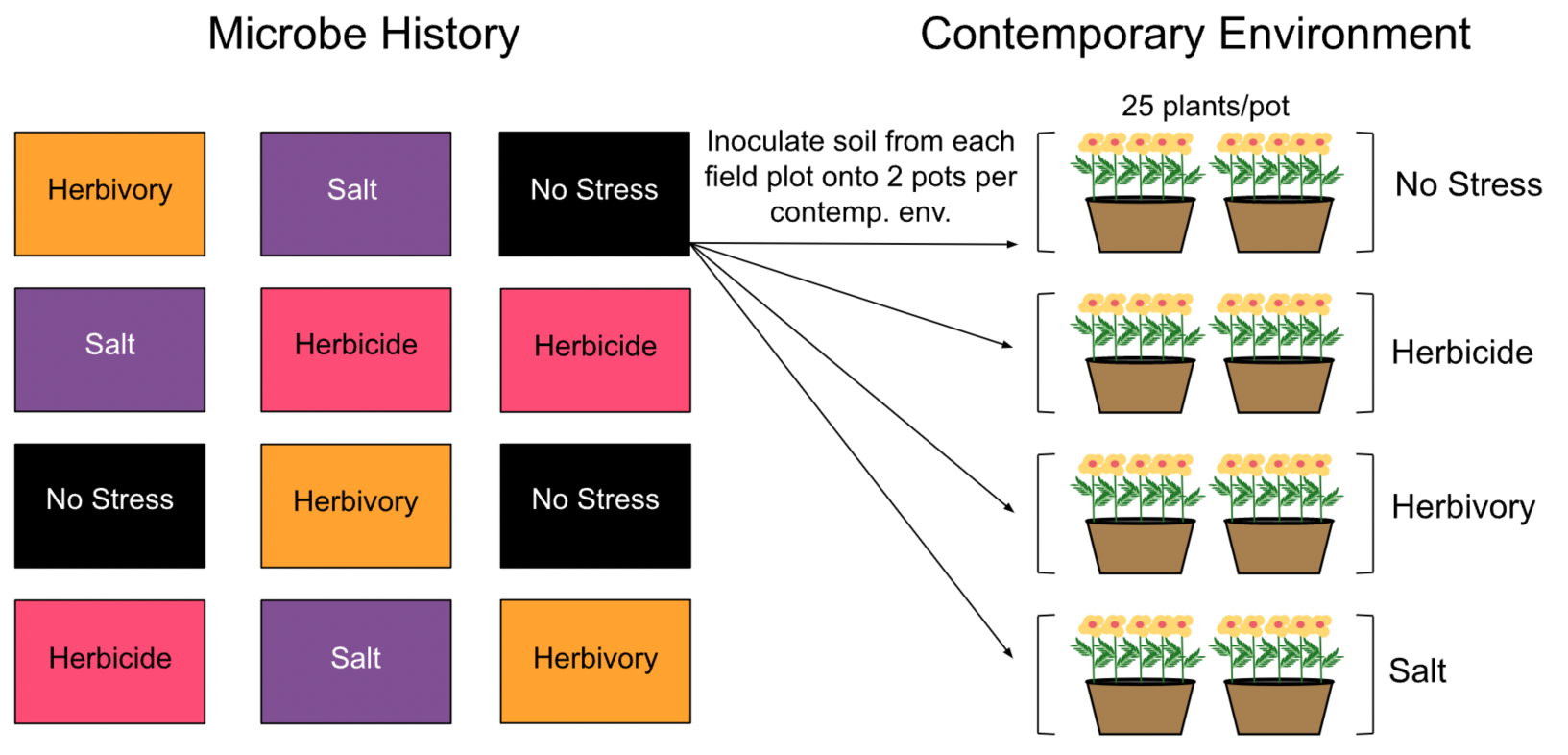
Experimental design. Each rectangle on the left represents a randomized field plot. ‘Microbe history’ is the treatment applied to each field plot, and rhizosphere soil containing microbes that had responded to the field treatments was taken from each field plot and inoculated into glasshouse mesocosms that were planted with new *Chamaecrista fasciculata* plants. ‘Contemporary environment’ is the treatment applied to experimental plants in glasshouse mesocosms. An individual from each of 50 full-sib families was planted into a random mesocosm location in each field plot × contemporary environment treatment. N = 2,400 plants; n = 96 mesocosms.

### Microbial Inocula from Field Experiment

For our microbial inocula, we capitalized on a field experiment that occurred during the summer of 2018 at the W.K. Kellogg Biological Station (KBS, Hickory Corners, Michigan, USA) in which a single individual from each of 100 *Chamaecrista* full-sib families was planted into each of twelve 2 × 2 m plots. To generate these full-sib families, seeds were collected in 2015 from maternal plants growing in two restored prairies in southwest Michigan (42°28′23′′N, 85°26′50′′W and 42°26′37′′N, 85°18′34′′W) that had been planted with identical prairie seed mixes in 2010 (Shooting Star Native Seeds; Houston County, MN). These field collected seeds were propagated in the glasshouse where controlled crosses were conducted (see Magnoli 2020 for details).

Field plots were treated with one of four stress treatments: salt, herbicide, simulated herbivory, or an unstressed control (n = 3 field plots per stress treatment). Salt plots were treated with 50 mL of 12 g L^-1^ of NaCl (Morton Salt, Chicago, IL, USA) 23 days after planting and with 50 mL of 24 g L^-1^ NaCL 38 and 45 days after planting, with salt solutions applied to the base of each plant each time. Foliage in herbicide plots was sprayed twice with glyphosate (0.02 and 0.03 kg active ingredient acre^-1^ applied 29 and 40 days after planting, respectively; these concentrations are ∼1/10 those applied to agricultural fields and were chosen based on pilot studies to stress plants without causing excess mortality). All fully expanded leaves in simulated herbivory plots were removed using scissors 23-27 days after planting and again 38 days after planting (these are extreme herbivory events that simulate the leaf removal that occurs with browsing or extreme insect herbivory). Note that effects of the herbivory treatment on soil microbial communities are mediated entirely by the plant, while effects of the herbicide and salt treatments may be partially mediated by the plant and partially due to the herbicide and salt interacting directly with the soil.

At the end of the growing season, after all fruits had matured, we collected 3 liters of rhizosphere soil from each field plot by shaking soil off the roots of *Chamaecrista* plants that were harvested from that plot (soils were stored in plastic baggies at 4 °C for three weeks before planting the glasshouse experiment). We removed plant material and macrobes (e.g., earthworms), then used the soil from each plot as inoculum in our glasshouse experiment.

### Glasshouse Experiment

We surface-sterilized 5-gallon pots (Zarn Inc 2000x, Reidsville, NC, USA) in 0.5% Physan 20 (Maril Products, Inc., Tustin, CA) and filled them with a sterilized base soil (autoclaved at 80 °C for two 4-h periods with a 48-h resting period in between) composed of a 7:3 mixture of Metro-Mix 360 (Sun Gro Horticulture, Bellevue, WA, USA) and sand (Quickrete All Purpose Sand, Atlanta, Georgia, USA). We then inoculated each mesocosm with a 200 mL layer of live field soil inoculum (1% live soil by volume) and topped each with a thin layer of sterile base soil to reduce contamination between mesocosms. We planted 50 full-sibling families of *Chamaecrista* into these mesocosms (one individual of each family into each field plot × contemporary environment treatment). These full-sib families were from the same crossing design as described in the “microbial inocula from field experiment” section above, and thus did not experience salt, herbicide, or herbivory stress. We scarified and imbibed seeds by nicking off a corner of the seed coat with a razor blade and submerging seeds in DI water for three days before planting into inoculated mesocosms. Some seeds did not emerge, so were re-planted 11 days after the original planting; however, flowering was significantly delayed for these plants, so we excluded them from analyses (n = 109 plants). Plants were maintained at 30 °C : 18 °C, day : night temps on a 15 hr : 9 hr, light : dark cycle (light was supplemented with 1000-W high pressure sodium lights) to mimic growing season conditions and were watered as needed (generally once daily). We manually removed weeds weekly that germinated from the live soil inoculum.

We applied stress treatments similar to those imposed in the field experiment five weeks after planting (modifications to the stress treatments were intended to achieve a dose that would stress plants without causing excess mortality based on outcomes from the field experiment). For the salt stress treatment, we applied 500 mL of a 12 g L^-1^ concentration of NaCl evenly across each mesocosm. For the herbicide stress treatment, we sprayed ∼800 mL of Roundup (Monsanto, Anvers, Belgium) at a concentration of 0.05 kg active ingredient (glyphosate) per hectare evenly across plants so that each leaf was nearly dripping (this concentration is 6-9% of field recommendations, so likely mimics pesticide drift rather than direct application). For the herbivory stress treatment, we removed all leaves more than 2.5 cm long using scissors seven weeks after planting to simulate a severe herbivory event. Clipping simulates herbivory in the first half of the growing season, and plants appeared to fully compensate by the end of the growing season as clipping did not significantly reduce final plant biomass (Tukey contrast: *P* = 0.15; Bolin, 2023). Each stress treatment was applied to all microbe history treatments (a 4 × 4 factorial design) resulting in 96 mesocosms (n = 2 mesocosms per field plot × 3 field plots/microbe history × 4 microbe histories × 4 contemporary environments = 96 mesocosms).

We measured two plant traits that are known to respond to both stress and microbes, and that are genetically variable in this population (Bolin, 2023): flowering time and specific leaf area (SLA; leaf area/leaf dry mass) (Lau & Lennon, 2011, 2012; Wagner *et al*., 2014; Panke-Buisse *et al*., 2015). We recorded flowering date for all plants. However, we only measured SLA for plants that were salt-stressed or unstressed, and inoculated with the microbes from the salt or unstressed history (N = 600 plants). We did this because it was infeasible to measure SLA on all plants, and because past studies suggest that salt may be the most relevant of our stressors to selection on SLA given that salt causes desiccation stress in plants and plants commonly reduce SLA in response to desiccation stress (Wright *et al*., 2001; Ackerly, 2004). We measured SLA 8.5 weeks after planting. After 15.5 weeks we harvested, dried (for at least two weeks at 60°C), and weighed all shoot biomass.

### Statistical Analyses

#### How do stress and microbial responses to stress influence natural selection on plant traits?

To estimate the strength and direction of natural selection on measured traits, we conducted genotypic selection analyses which regress relative fitness onto standardized trait values (Rausher, 1992; see also Lande & Arnold, 1983). Genotypic selection analyses use family mean fitness and trait values, which removes biases due to environment-induced covariances between traits and fitness (Rausher, 1992) and reduces potential biases due to the invisible fraction (i.e., individuals that die before a trait is measured or expressed; Hadfield, 2008), although these biases are not completely eliminated because siblings are not genetically identical. We standardized traits (SLA and flowering time) globally by subtracting the global mean trait value from each observation and dividing by the global standard deviation of trait values (De Lisle & Svensson, 2017). We also conducted additional analyses accounting for variation in productivity among mesocosms to test whether stress and microbe history directly altered plant natural selection, or whether they affected selection indirectly by altering the competitive environment (e.g., by increasing mortality). In no cases were effects of microbes on plant selection mediated by changes in the competitive environment, so we present these statistical methods and results in the Supporting Information (Methods S1; Tables S1-S7; Fig. S1). All statistical analyses were conducted in R version 4.2.0 (R Core Team, 2018).

*Viability selection –* To test whether contemporary stress and/or microbe history affect viability selection gradients, which measure direct selection acting on a trait after accounting for selection acting on measured correlated traits, we fit weighted binomial generalized linear models (used when the dependent variable is a proportion) with a logit link function. We included probability of survival to flower (proportion of surviving individuals in each family) as the response variable; a plant that did not survive to flower had zero fitness, while a plant that did survive to flower likely had nonzero male fitness at minimum, as well as potentially some female fitness. We included standardized trait values (flowering time and SLA), microbe history, contemporary environment, and all interactions as predictors. We weighted each data point by the number of individuals in each family × treatment combination (range: 1-3 due to some seedlings failing to emerge; mode: 3). For selection gradients, we were limited to the contemporary unstressed and salt stressed treatments with microbes from unstressed and salt stressed field plots because we only measured multiple traits (SLA and flowering time) in these treatments. Significant trait × cotemporary environment interactions indicate that stress affects natural selection on plant traits; while significant trait × microbe history interactions indicate that microbial responses to stress alter natural selection. To test for significant differences among microbe history and contemporary stress treatments we performed post-hoc Tukey tests using the *emtrends* function in the ‘emmeans’ package (Lenth *et al*., 2018).We estimated linear selection gradients by fitting a separate model for each microbe history × contemporary environment treatment that included probability of survival to flower as the response variable, and both standardized trait values as predictors. We then transformed these logistic coefficients into coefficients that can be used in microevolutionary equations to predict evolutionary change (*β_avggrad_*) following the methods of Janzen & Stern (1998).

To test whether contemporary stress and/or microbe history affect selection differentials, which measure total selection acting on a trait including both direct and indirect selection due to selection acting on correlated traits, we fit similar models as described in the previous paragraph except we fit a separate model for each plant trait (i.e., only a single trait was included in each model). As above, we estimated linear selection differentials by fitting a separate model for each microbe history × contemporary environment treatment. For SLA we estimated differentials in the same subset of treatments that we used to estimate gradients (contemporary unstressed and salt stressed treatments with microbes from unstressed and salt stressed field plots), while for flowering time we estimated differentials in all contemporary environment and microbe treatments. We transformed these logistic coefficients and tested for significant differences among treatments as described for selection gradients above.

For all models, we also tested for nonlinear viability selection by including quadratic terms for each trait and interactions between quadratic terms and the contemporary environment and microbe history treatments. Nonlinear selection coefficients were doubled as in Stinchcombe *et al*. (2008). Because nonlinear selection was rarely statistically significant, we focus on linear selection in the main text (although we present quadratic selection coefficients in Tables 1 and 2), but present results from models testing whether the contemporary environment and microbe history affect nonlinear selection in Tables S5-S7.

**Table 1.** Linear and quadratic genotypic **(a)** viability and **(b)** fecundity selection differentials. Coefficients were estimated from linear regressions of relative fitness on standardized trait values (Lande & Arnold, 1983). Quadratic coefficients were doubled as in Stinchcombe *et al*. (2008). Standard errors are shown in parentheses. For viability differentials, logistic coefficients were transformed according to the methods of Janzen & Stern (1998). Linear coefficients were estimated from models containing only linear terms, while quadratic coefficients were estimated from the full models. † *P* < 0.1, * *P* < 0.05, ** *P* < 0.01, *** *P* < 0.001. ^a^ Unusually high quadratic viability differential and error are due to only one family having imperfect survival when grown in contemporary herbivory with salt microbes.

| Contemporary Environment | Microbe History | Flowering Time |  | SLA |  |
| --- | --- | --- | --- | --- | --- |
|  |  | Linear | Quadratic | Linear | Quadratic |
| (a) Viability differentials |  |  |  |  |  |
| No Stress | No Stress | 0.05 (0.05) | 0.14 (0.16) | -0.04 (0.06) | -0.01 (0.14) |
|  | Salt | -0.04 (0.04) | 0.12 (0.16) | -0.09 (0.05) † | -0.21 (0.11) |
|  | Herbicide | 0.08 (0.05) † | -0.05 (0.06) | --- | --- |
|  | Herbivory | -0.08 (0.05) | 0.02 (0.11) | --- | --- |
| Salt | No Stress | 0.13 (0.09) | -0.15 (0.09) † | **0.17 (0.08) \*** | **-0.18 (0.08) \*** |
|  | Salt | 0.03 (0.10) | -0.01 (0.29) | -0.04 (0.05) | **-0.12 (0.06) \*** |
|  | Herbicide | 0.05 (0.07) | -0.09 (0.11) | --- | --- |
|  | Herbivory | -0.05 (0.07) | -0.11 (0.09) | --- | --- |
| Herbicide | No Stress | 0.06 (0.06) | -0.02 (0.09) | --- | --- |
|  | Salt | -0.08 (0.04) † | -0.04 (0.05) | --- | --- |
|  | Herbicide | 0.02 (0.06) | 0.10 (0.09) | --- | --- |
|  | Herbivory | -0.06 (0.06) | -0.13 (0.13) | --- | --- |
| Herbivory | No Stress | -0.02 (0.05) | -0.15 (0.10) | --- | --- |
|  | Salt | 0.01 (0.02) | 42.34 (1434.58)<sup>a</sup> | --- | --- |
|  | Herbicide | -0.05 (0.03) | -0.03 (0.05) | --- | --- |
|  | Herbivory | 0 (0.06) | 0 (0.16) | --- | --- |
| (b) Fecundity differentials |  |  |  |  |  |
| No Stress | No Stress | **-0.53 (0.16) \*\*** | -0.16 (0.18) | **-0.72 (0.22) \*\*** | 0.22 (0.32) |
|  | Salt | -0.24 (0.15) | -0.56 (0.23) | **-0.72 (0.15) \*\*\*** | 0.28 (0.23) |
|  | Herbicide | **-0.65 (0.18) \*\*\*** | 0.20 (0.14) | --- | --- |
|  | Herbivory | **-0.36 (0.15) \*** | -0.14 (0.13) | --- | --- |
| Salt | No Stress | -0.12 (0.10) | -0.02 (0.06) | **-0.29 (0.12) \*** | 0.56 (0.14) † |
|  | Salt | -0.20 (0.13) | -0.12 (0.20) | **-0.20 (0.07) \*\*** | 0.16 (0.08) |
|  | Herbicide | -0.22 (0.22) | -0.10 (0.20) | --- | --- |
|  | Herbivory | **-0.23 (0.09) \*** | 0.02 (0.06) | --- | --- |
| Herbicide | No Stress | -0.16 (0.15) | -0.42 † (0.11) | --- | --- |
|  | Salt | **-0.31 (0.14) \*** | -0.06 (0.08) | --- | --- |
|  | Herbicide | **-0.33 (0.14) \*** | -0.18 (0.10) | --- | --- |
|  | Herbivory | -0.30 (0.18) | **-0.80 \* (0.19)** | --- | --- |
| Herbivory | No Stress | **-0.29 (0.11) \*** | -0.12 (0.10) | --- | --- |
|  | Salt | -0.15 (0.16) | **-0.94 \* (0.19)** | --- | --- |
|  | Herbicide | **-0.25 (0.12) \*** | -0.26 (0.09) | --- | --- |
|  | Herbivory | **-0.49 (0.18) \*** | 0.62 (0.26) | --- | --- |

**Table 2.** Linear and quadratic genotypic **(a)** viability and **(b)** fecundity selection gradients. Coefficients were estimated from linear regressions of relative fitness on standardized trait values (Lande & Arnold, 1983). Quadratic coefficients were doubled as in Stinchcombe *et al*. (2008). Standard errors are shown in parentheses. For viability differentials, logistic coefficients were transformed according to the methods of Janzen & Stern (1998). For selection gradients, we were limited to the contemporary no stress and salt stress treatments with microbes from unstressed and salt stressed field plots because we only measured SLA in these treatments. Linear coefficients were estimated from models containing only linear terms, while quadratic coefficients were estimated from the full models. † *P* < 0.1, * *P* < 0.05, ** *P* < 0.01, *** *P* < 0.001.

| Contemporary Environment | Microbe History | Flowering Time |  | SLA |  |
| --- | --- | --- | --- | --- | --- |
|  |  | Linear | Quadratic | Linear | Quadratic |
| (a) Viability gradients |  |  |  |  |  |
| No Stress | No Stress | 0.06 (0.05) | 0.14 (0.16) | -0.05 (0.05) | -0.04 (0.12) |
|  | Salt | -0.03 (0.04) | 0.10 (0.15) | -0.06 (0.05) | -0.19 (0.10) † |
| Salt | No Stress | 0.14 (0.09) | -0.15 (0.09) † | **0.15 (0.07) \*** | -0.14 (0.08) † |
|  | Salt | 0 (0.1) | -0.09 (0.30) | 0.07 (0.07) | -0.06 (0.09) |
| (b) Fecundity gradients |  |  |  |  |  |
| No Stress | No Stress | **-0.38 (0.17) \*** | -0.20 (0.18) | **-0.57 (0.22) \*** | 0 (0.31) |
|  | Salt | -0.16 (0.13) | -0.20 (0.21) | **-0.67 (0.16) \*\*\*** | 0.28 (0.24) |
| Salt | No Stress | -0.08 (0.09) | 0 (0.05) | **-0.33 (0.11) \*\*** | 0.52 (0.14) † |
|  | Salt | -0.11 (0.13) | -0.02 (0.19) | **-0.19 (0.08) \*** | 0.20 (0.09) |

*Fecundity selection –* To test whether contemporary stress and/or microbe history affect fecundity selection gradients and differentials, for all surviving individuals we fit linear models as described for viability selection analyses except we included ln-transformed aboveground biomass (family mean) as the response variable. We used aboveground biomass as a proxy for seedset because it correlates strongly and positively with fruit production in *Chamaecrista* (Galloway & Fenster, 2001) and because it was infeasible to hand pollinate the thousands of plants included in this experiment. To estimate the strength and direction of selection in each treatment, we calculated linear selection gradients and differentials as described for survival above, but with raw relative aboveground biomass (non-transformed family mean) as the response variable. We used raw biomass to estimate selection coefficients because transformed fitness can invalidate the interpretation of selection analyses and lead to inaccurate estimates of selection (Lande & Arnold, 1983; Mitchell-Olds & Shaw, 1987), but we used ln-transformed biomass in hypothesis testing (i.e., testing whether microbe history or contemporary environment affects selection) to better meet model assumptions. We relativized biomass globally by dividing by global mean fitness (as opposed to within-treatment mean fitness), which is recommended for traits that are expected to undergo hard selection (i.e., whose adaptive value is independent of density; De Lisle & Svensson, 2017).

#### How do stress and microbial responses to stress influence plant opportunity for selection?

To test whether stress and microbial responses to stress influenced plant opportunity for selection (*I*, equal to the variance in relative fitness), we first calculated *I* as the variance in relative biomass for each mesocosm. We then fit a linear model that included *I* as the response variable, and microbe history, contemporary environment, and their interaction as fixed effects. To test for significant differences among stress and microbe history treatments, we performed post-hoc Tukey tests using the ‘emmeans’ package (Lenth *et al*., 2018).

The variance in relative fitness is equal to the ratio of variance in absolute fitness to squared mean absolute fitness (Wade & Shuster, 2005). Therefore, to test whether changes in *I* were due to changes in mean fitness vs. changes in the variance in fitness, we calculated each of these for each mesocosm and conducted similar analyses to those described in the previous paragraph (Waterton *et. al.*, 2022).

### Quantifying drivers of selection

To quantify the effect of stress itself, the effect of microbial community responses to stress, and the legacy effect of microbial responses to stress on plant selection (i.e., to quantify the arrows depicted in Fig. **1a-d**, as shown in Fig. **1e-h**), we used the following equations for each stress environment where *β* is the estimated selection coefficient (differential or gradient), the first subscript is the contemporary environment, and the second subscript is the microbe history:

1. Effect of stress on selection = *β _Stress, No Stress_ – β _No Stress, No Stress_*
2. Effect of microbial response to stress on selection = *β _Stress, Stress_ – β _Stress, No Stress_*
3. Legacy effect of microbial responses to stress on selection = *β _No Stress, Stress_ – β _No Stress, No Stress_*

To test whether microbial responses to stress typically opposed or reinforced the effects of stress itself, we conducted a sign test using the binom.test function. Note that the legacy effect of microbial responses to stress on selection (Equation 3) simulates a microbial legacy of stress for the next generation of annual plants after the stress ceases, as opposed to the generation of plants that experienced the stress. Additionally, estimated direct effects of stress on selection may include some effects that result from very rapid changes in microbial communities in response to the contemporary stress environment imposed during our glasshouse experiment.

We also quantified the effects of stress, microbial responses to stress, and microbial legacy effects of stress on plant selection gradients under salt stress (we were limited to salt stress for selection gradients, see ‘Glasshouse Experiment’ Methods). Results were similar for selection gradients and differentials, so we present selection gradient results in Fig. S2.

## RESULTS

### How do stress and microbial responses to stress influence natural selection on plant traits?

Salt stress weakened fecundity selection favoring lower SLA (i.e., caused the selection differential and gradient to be closer to zero) (fecundity differential, SLA × contemp. env.: *F*_1,173_ = 5.3, *P* = 0.023; fecundity gradient, SLA × contemp. env.: *F*_1,161_ = 2.8, *P* = 0.097; Fig. **3c**; Tables **1b**, **2b**; Tables S2, S4; Fig. S3) and reversed the direction of viability selection so that higher SLA (thinner leaves) was favored under salt stress (viability differential, SLA × contemp. env.: χ_1_^2^ = 4.4, *P* = 0.036; viability gradient, SLA × contemp. env.: χ_1_^2^ = 4.7, *P* = 0.030; Fig. **3a**; Table **1a**; Table **2a**; Tables S2, S4; Fig. S4). However, microbial responses to salt (i.e., microbes from the field salt treatment) counteracted the effect of salt stress on selection and tended to favor lower SLA (viability differential, SLA × microbe history: χ_1_^2^ = 6.2, *P* = 0.013; Fig. **3a**; Table **1a**; Table S2).

**Figure 3.**
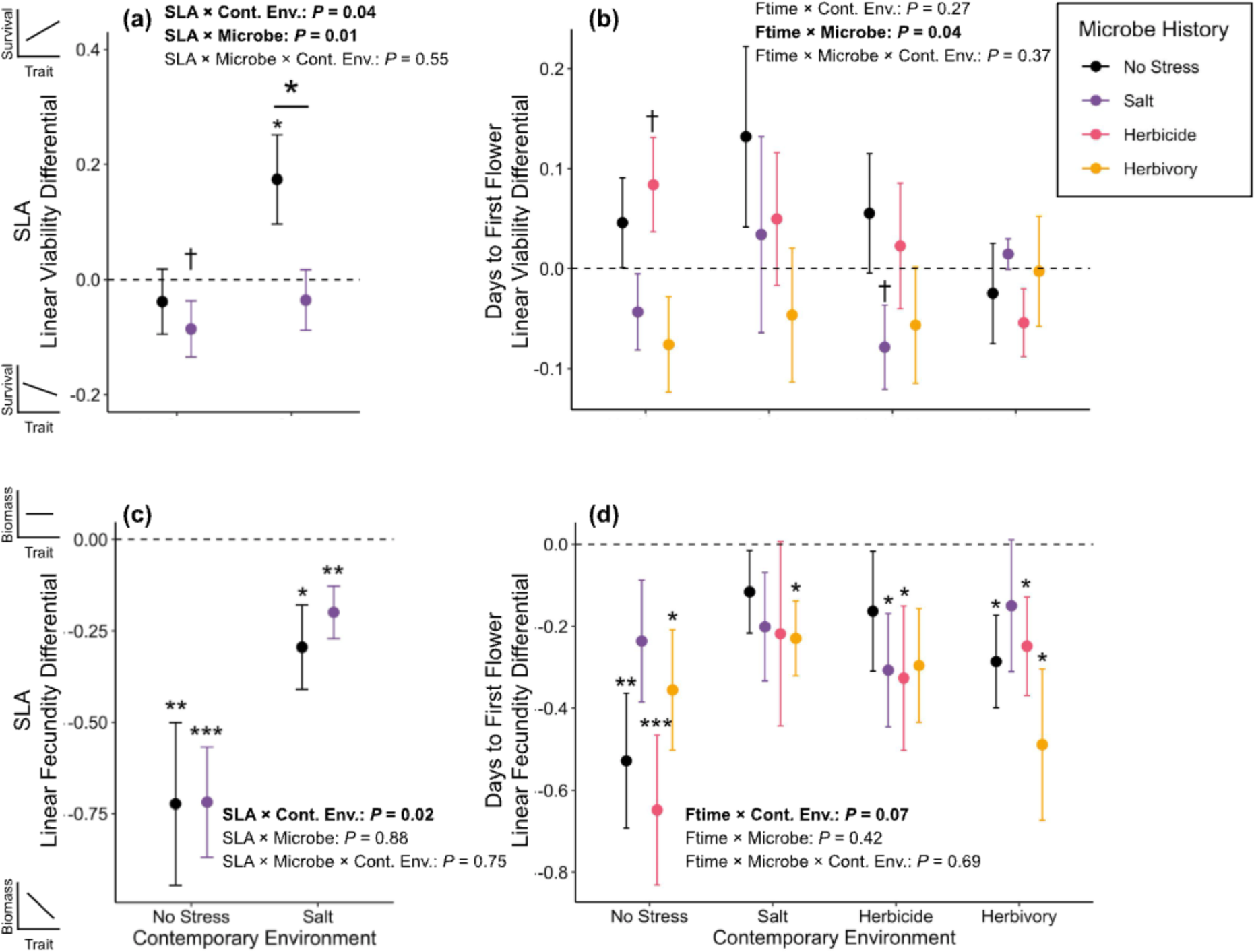
Stress and microbial responses to stress (‘microbe history’) affected viability selection **(a, b)** and fecundity selection **(c, d)** on plant specific leaf area (SLA; left column) and days to first flower (right column) in *Chamaecrista fasciculata*. Viability differentials represent the slope estimate of the trait-survival probability relationship in each microbe history × contemporary environment treatment, and fecundity differentials represent the slope estimate of the trait-biomass relationship (as depicted in the cartoon graphs on the y-axis in panels **a** and **c**). Differential estimates are from models that only include linear terms. Values above the dashed line indicate positive selection on the plant trait; values below the dashed line indicate negative selection. Error bars are fitted SE around the selection differential estimate. ‘No stress,’ ‘salt’, ‘herbicide,’ and ‘herbivory’ microbe history treatments represent soil microbial communities from field plots where plants were unstressed (black), salt-stressed (purple), herbicide-stressed (pink), and herbivory-stressed (yellow), respectively. Asterisk above bar indicates significant differences between microbe history treatments; asterisk/dagger above treatment indicates that selection is significantly different from zero; † *P* < 0.1, * *P* < 0.05, ** *P* < 0.01, *** *P* < 0.001.

Although viability selection on plant flowering time was typically weak and non-significant, microbe history significantly affected this selection such that microbial responses to herbivory tended to cause viability selection to favor earlier flowering relative to microbes from non-stressful environments and herbicide microbes, although these pairwise contrasts were not statistically significant (viability differential, flowering time × microbe history: χ_3_^2^ = 8.2, *P* = 0.041; Fig. **3b**; Tables **1a**, **2a**: Fig. S5; Table S3; Tukey contrasts: *P* = 0.15 and *P* = 0.27, respectively). Fecundity selection favored earlier flowering in all environments, and stress tended to weaken this selection (fecundity differential, flowering time: *P* < 0.01 in all environments, flowering time × contemp. env.: F_3,9_ = 2.4, *P* = 0.067; Fig. **3d**; Table S3; Fig. S6).

### Quantifying drivers of selection

Qualitative comparisons of effect sizes suggest that the effect of stress itself typically affected selection on plant traits more strongly than microbial community responses to stress (Fig. 4). However, microbial responses to stress often substantially affected selection, and in two cases the change in selection caused by microbial responses to stress exceeded effects of stress itself (flowering time viability selection under salt and herbicide stress). Microbial responses to stress opposed the effects of stress on selection in all cases but one (fecundity selection for higher SLA under salt stress), and this trend was marginally significant despite low statistical power (sign test: *P* = 0.070), suggesting that changes to microbial communities typically reduced the effects of stress on natural selection in our study (Fig. **4a**; Fig. **1b,c**). Furthermore, legacy effects of microbial responses to stress on plant flowering time selection, which occur when stress-adapted microbes influence selection in non-stressful environments, were generally quite strong. In fact, microbial legacy effects of stress were stronger than the effects of stress itself on flowering time viability selection in all stress environments (Fig. **4b**, Fig. **1d**). These legacy effects indicate that stress can continue to alter selection even after the stress has subsided because of lingering changes to microbial community composition.

**Figure 4.**
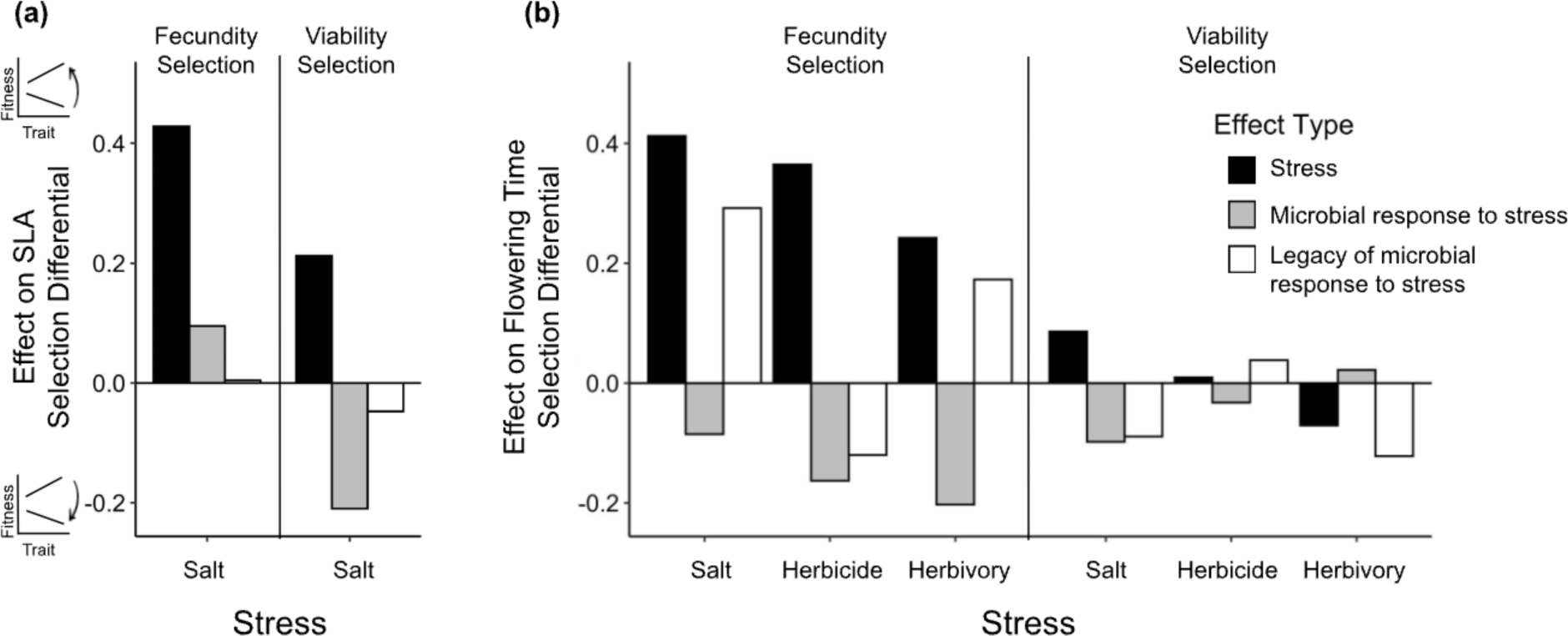
Selection on *Chamaecrista fasciculata* **(a)** SLA and **(b)** flowering time can be affected by the stress environment itself (black bars), by microbial community responses to stress (gray bars), or by microbial legacy effects of past stress (white bars). Legacy effects of microbial responses to stress occur when stress selects for microbial communities that alter selection even after the stress has ceased (i.e., in non-stressful contemporary environments). Effects of stress, microbial response to stress, and legacy of microbial response to stress were calculated as described in equations 1, 2, and 3. Values above zero indicate that an effect caused selection differentials to become more positive (or less negative), and values below zero indicate an effect caused selection to become more negative (or less positive), as depicted in the cartoon graphs on the y-axis in (a). Fecundity and viability selection are shown on the left and right side of each panel, respectively, and contemporary stress environments are shown across the x-axis.

### How do stress and microbial responses to stress influence plant opportunity for selection?

Herbicide stress affected the opportunity for selection (the variance in relative fitness; *I*) and also caused changes to the soil microbial community that affected *I*. Herbicide stress itself increased *I* relative to non-stressful environments and herbivory stress (Tukey contrasts: *P* = 0.003 and *P* = 0.015, respectively) and tended to increase *I* relative to salt stress (Tukey contrast: *P* = 0.085) (contemp. env.: *F*_3,80_ = 5.2, *P* = 0.003; Fig. **5a**; Table S8). Microbial communities from the herbicide treatment also increased *I* relative to microbial communities from salt and herbivory treatments (Tukey contrasts: *P* < 0.001 and *P* = 0.027, respectively) (microbe history: *F*_3,80_ = 6.4, *P* < 0.001; Fig. **5a**; Table S8). Additionally, microbes from the no-stress treatment increased *I* relative to microbes from the salt treatment (*P* = 0.047).

**Figure 5.**
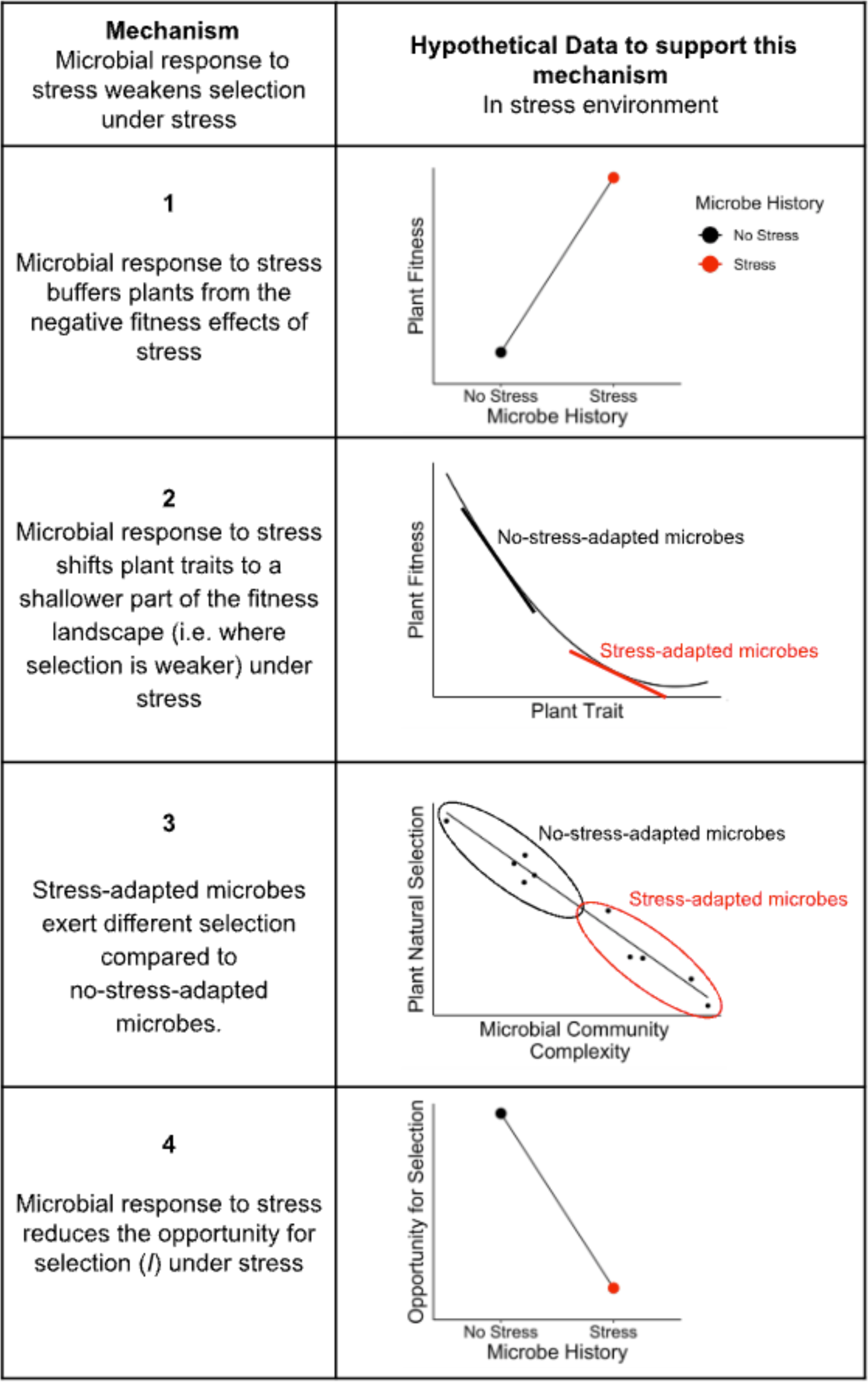
(a) Stress and microbial community responses to stress (microbe history) both influenced plant opportunity for selection (*I*), which limits the maximum strength of selection that can act on a population. The opportunity for selection is equal to the variance in relative fitness, or equivalently, *σ^2^_W_*/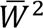, and **(b)** microbial responses to stress altered *I* by altering both mean absolute fitness (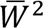) and variance in absolute fitness (*σ^2^_W_*) in *Chamaecrista fasciculata*. No stress, salt, herbicide, and herbivory microbe history treatments represent soil microbes from field plots where plants were unstressed (black), salt-stressed (purple), herbicide-stressed (pink), and herbivory-stressed (yellow), respectively. In **(b)**, contemporary no stress, salt, herbicide, and herbivory environments are shown as squares, diamonds, circles, and triangles, respectively. The dashed line is the identity line where *I* = 1, and treatments in the lower right and upper left have *I* < 1 and *I* > 1, respectively. Error bars are SE.

The opportunity for selection is determined by both the mean (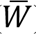) and variance (*σ^2^_W_*) in absolute fitness (*σ^2^_W_*/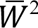; Wade & Shuster, 2005), so microbes and/or stress can alter *I* by changing plant 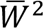 (Fig. **5b** x-axis) or by changing plant *σ^2^_W_* (Fig. **5b** y-axis). Greater *I* under contemporary herbicide stress compared to non-stressful environments was driven by herbicide reducing 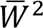 (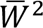, contemp. env.: *F_3,80_* = 22.4, *P* < 0.001; Fig. **5b**; Table S8; Tukey contrast: *P* < 0.001). Greater *I* under contemporary herbicide stress compared to salt stress resulted because, while herbicide stress increased both 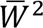and *σ^2^_W_* compared to salt stress (*σ^2^_W_*, contemp. env.: *F_3,80_* = 6.3, *P* < 0.001; Fig. **5b**; Table S8; Tukey contrasts: *P* = 0.050 and *P* = 0.006, respectively), it increased *σ^2^_W_* proportionally more. By contrast, microbial responses to stress did not significantly affect 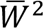or *σ^2^_W_,* but had small effects on both that cumulatively resulted in significant effects on *I* (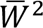, microbe history: *F_3,80_* = 1.6, *P* = 0.20; *σ^2^_W_*, microbe history: *F_3,80_* = 0.6, *P* = 0.64; Fig. **5b**; Table S8).

## DISCUSSION

When environments change, plants encounter not only new abiotic conditions and aboveground biotic interactions, but also potentially new belowground interactions with soil microbial communities that have responded to these environments. By independently manipulating the stress environment and the soil microbes selected for in that stress environment, we showed that microbial responses to stress affect plant natural selection. While the effects of microbial community responses were generally weaker than the effects of stress itself, they substantially modified the strength, and occasionally the direction, of selection. Our findings also suggest that these microbial responses to stress may lead to legacy effects of stress; in other words, the effects of stress on selection may persist even after the stressful conditions have ceased because of lasting changes to the soil microbial community.

### Community context affects plant evolutionary responses to stress

Community context may commonly affect evolution in response to stress and other global change factors (Lau & terHorst, 2020). For example, elevated CO_2_ only altered natural selection on plant growth traits in the presence of competitors (Lau *et al*., 2014), and warming only accelerated the evolution of *Daphnia* life history traits in the presence of a predator (Tseng & O’Connor). Here, we showed that natural selection acting on plants is affected not only by the presence of other community members, but in this case also by the particular assembly of microbes plants are likely to encounter in various stressful environments.

We found that microbial community responses to stress counteracted the selective effects of stress itself in every case but one (fecundity selection on plant SLA under salt stress). As a result, the responses of microbial communities to stress weakened the effects of stress on natural selection. In other words, in cases where the effect of stress was to strengthen natural selection (e.g., viability selection on plant SLA), stress-adapted microbes (i.e., microbes from that stress treatment in the field) exerted selection in the opposite direction, whereas in cases where the effect of stress was to weaken natural selection (e.g., fecundity selection on plant flowering time), stress-adapted microbes strengthened natural selection. Similar to how community diversity can promote plant population stability (Tilman & Downing, 1994), these patterns suggest that microbial responses to stress may promote plant evolutionary stasis by buffering plants from extreme swings in the strength of natural selection when the environment shifts.

Moreover, microbial responses to stress also had relatively strong legacy effects on selection in non-stressful environments, particularly for flowering time. As a result, stress-induced changes to microbial communities may have lingering effects on plant evolution for the next generation of plants, even when those plants do not experience stress themselves. Microbial legacy effects were equally likely to oppose the effects of stress itself as they were to reinforce them, suggesting that microbial legacies may sometimes reverse the evolutionary effects of stress and other times extend them. Legacy effects of stress on ecological variables are not uncommon, and in some cases these legacy effects of stress occur via longer-lasting effects on microbial communities. For example, drought stress can change soil microbes in ways that alter plant competitive outcomes in future generations (Kaisermann *et al*., 2017), and in fact a microbial legacy of drought may affect plants even more strongly than a plant legacy (i.e., maternal effects) of drought (De Long *et al*. 2019). Our results suggest that these microbial legacy effects also can affect plant evolutionary processes.

Surprisingly, microbial legacy effects of stress were equally likely to oppose vs. reinforce effects of microbial responses to stress. One might expect legacy effects to parallel (but perhaps be weaker than) microbial response effects if they result from persistent changes to the microbial community. In cases where legacy effects oppose microbial response effects, it is likely that stress-adapted microbial communities differentially affect plant natural selection depending on the contemporary environment. In other words, the microbial response to stress might increase (or decrease) the strength of selection under stress, but have the opposite effect in a non-stressful environment. Such environment-specific effects of microbes are not uncommon (e.g., nitrogen fertilization reduces plant response to mycorrhizal fungi; Hoeksema *et al*. 2010). Alternatively, microbial communities may have shifted to new community states post stress (distinct from the stress-adapted community and the no-stress-adapted community). Tracking microbial community composition through time might help differentiate between these possibilities.

### Microbial responses to stress alter the strength of natural selection on plants: Potential mechanisms

We found that stress-adapted soil microbes altered the strength of natural selection on plant traits, and there are several mechanisms by which this could happen. Stress-adapted microbes could alter natural selection by affecting the opportunity for selection in plants (see next section; Table 3, Mechanism 4), but also by altering the fitness effects of stress, by shifting plants to a steeper or shallower part of the fitness landscape, or simply if microbial communities are selective agents on plant traits and the strength of selection depends on microbial community properties (Table 3). Here we focus on how stress-adapted microbes could weaken natural selection under stress because this was the most common pattern in our study, but these same mechanisms (with the reverse direction of effect) could also contribute to microbes strengthening selection on plant traits. We focus on how microbes could affect selection on plant stress tolerance/avoidance traits (such as flowering time and SLA), but microbes could also alter selection on other traits, including traits that mediate plant-microbe interactions, via these mechanisms (Angulo *et al*., 2022).

**Table 3.**
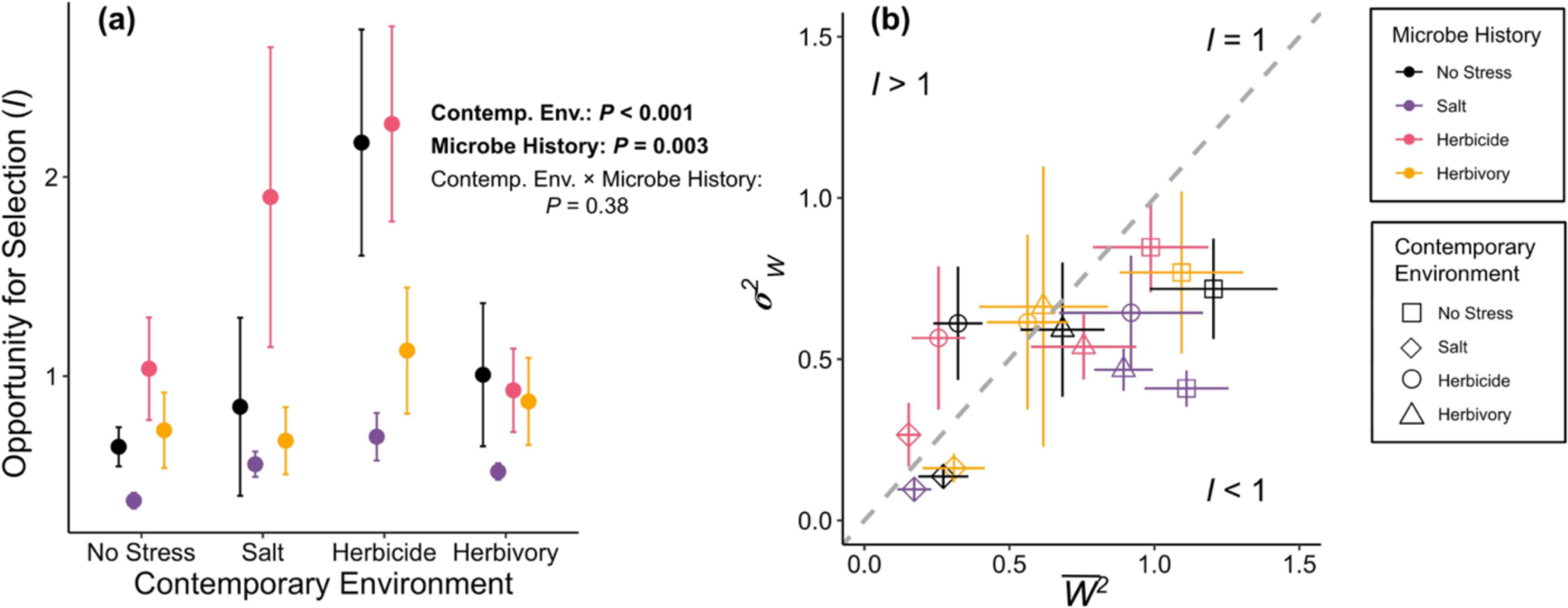
Potential mechanisms by which soil microbial community responses to stress could influence natural selection on plant stress tolerance/avoidance traits. *Hypothetical data show stress-adapted microbes weakening natural selection on plant traits as was most commonly found in our study, but note that support from our study for mechanisms 2 and 3 come from cases where microbial responses to stress strengthened, rather than weakened, plant natural selection.

First, stress-adapted microbes could weaken plant natural selection under stress if they buffer plants from the negative fitness effects of stress, as is commonly found (Petipas *et al*., 2021; Table 3, Mechanism 1). For example, biofilm-producing microbes are often favored in low soil moisture conditions, and microbial biofilm production protects plants from drought stress (Bolin *et al*. 2022). If plants are fully protected from stress by the microbial community that develops in that stress environment, then there may be little fitness advantage to stress-tolerant plant phenotypes. This mechanism could have contributed to weaker viability selection on plant SLA with salt microbes under salt stress because salt microbes tended to reduce the negative effects of salt on plant survival and aboveground biomass (Bolin, 2023).

Second, stress-adapted microbes could weaken (or strengthen) plant natural selection under stress if there is curvature in the fitness landscape (i.e., the selection differential or gradient is non-linear), and stress-adapted microbes alter plant traits in ways that shift the population to a shallower (or steeper) part of the fitness landscape (Table 3, Mechanism 2). Microbes commonly affect plant traits (reviewed in: Friesen *et al*., 2011; Goh *et al*., 2013), so when microbial communities shift in ways that alter mean plant trait values it could result in changes in the strength of selection without any changes in the trait-fitness relationship. In our study, herbivory microbes strengthened fecundity selection on plant flowering time and tended to accelerate flowering relative to microbes from non-stressful environments (Bolin, 2023). The flowering time fitness landscape had positive curvature, meaning this microbe-mediated trait shift would move the population to a steeper part of the curve (Fig. S5d), suggesting that this mechanism could have contributed to microbial effects on plant selection in our study.

Third, microbial communities themselves can be selective agents on plant traits (Lau & Lennon, 2011; Wagner *et al*., 2014; Chaney & Baucom, 2020). When stress alters microbial community composition, then stress-adapted microbial communities could weaken plant natural selection under stress if the selection imposed by the stress-adapted microbial community is in the opposing direction as the effect of stress (Table 3, Mechanism 3). For example, stress might reduce microbial diversity (Lozupone & Knight, 2007), and simpler, less diverse soil microbial communities can exert stronger selection for earlier flowering than more complex microbial communities (Lau & Lennon, 2011; but see Chaney & Baucom, 2020). If stress reduces microbial complexity and reduced microbial complexity also strengthens selection for earlier flowering in our system, then this pattern could explain the tendency for plants inoculated with herbicide and salt microbes to experience stronger selection for earlier flowering, although we cannot determine to what extent this is due to this mechanism vs. the preceding two mechanisms.

### Stress and microbial responses to stress affect plant opportunity for selection

Stress affected plant opportunity for selection (*I*) both directly and by changing the soil microbial community, suggesting that variation in soil microbial community composition can contribute to the maximum strength of selection experienced by plants, and therefore the maximum amount of phenotypic change that can occur in a generation (Wade & Shuster, 2005). Herbicide stress itself increased *I* and shifted microbial community composition in a way that also increased *I*, indicating that herbicide-stressed plant populations have the potential to undergo strong selection (due in part to soil microbial community responses to stress). Although the traits we measured in our study did not experience particularly strong selection under herbicide stress, other unmeasured traits may have been under strong selection. For example, evolution of herbicide resistance often occurs via mutation at the site of herbicide action (i.e., enzymes or proteins where herbicides bind and disrupt plant function) or the ability to metabolize herbicide (Prather *et al*., 2000). By contrast, microbial responses to salt generally reduced *I* which could slow plant evolution under salt stress (Arnold & Wade, 1984; Crow, 1958; Caruso *et al*., 2017; Table 3, Mechanism 4). In general, our findings suggest that different stressors may influence plant *I* not only directly, but also by altering the soil microbial community, with some microbial communities increasing *I* and others decreasing *I*.

Ultimately, the outcomes of microbe-driven changes in *I* for plant populations could be positive or negative. For example, greater *I* (e.g., with herbicide microbes) and the resulting stronger selection could lead to greater phenotypic change that could help a population quickly adapt to a stressful environment. However, this stronger selection may also come at the cost of a higher demographic toll that increases the chance of extinction (the ‘cost of selection’; Haldane, 1957). Likewise, lower *I* (e.g., with salt microbes) and the resulting weaker selection could hinder plant adaptation to stressful environments but spare populations from a high cost of selection. Or, populations may have low mean fitness and very low variance in fitness, potentially resulting in both low *I* and a high demographic toll. In general, the advantageousness of microbe-driven changes in plant *I* will depend on the extent to which beneficial phenotypic changes are limited by variance in relative fitness in a population, and the extent to which raising the ceiling of maximum selection strength puts populations at demographic risk.

### Implications

We showed that stress can alter patterns of natural selection by changing soil microbial community composition, and that microbial legacies of stress can alter patterns of selection for the next generation of annual plants who themselves do not experience stress. However, these effects are likely to be sensitive to the strength and duration of stress events, and how quickly microbes respond to those stress events. For example, microbes may not respond sufficiently to a brief and relatively benign stress event to influence plant selection, or may respond too quickly to the lifting of stress to have meaningful legacy effects on natural selection for the next generation of plants. Alternatively, microbial responses to stress may be so rapid that effects of microbial community change cannot be partitioned from the effects of stress itself (e.g., Mackelprang *et al*. 2011). For example, even though our microbial history treatments persisted long enough to influence selection in our study, the effects of stress, microbial responses to stress, and microbial legacy effects of stress that we reported here all likely include additional microbe-mediated effects that occurred as our inoculated microbial communities rapidly responded to our glasshouse treatments. In our study, however, the effects of microbial responses to stress that we report are likely underestimated relative to the effects of stress itself because the effects of microbial responses to stress opposed the effects of stress on selection.

Despite these potential constraints and caveats, the microbial communities we used as inoculum responded to stress over the course of one growing season in the field (although it is also possible that they responded much more quickly), and we were able to detect legacy effects of those communities on a subsequent generation of plants in the glasshouse. Together these findings suggest that significant changes to microbial communities that in turn affect plant evolution can both occur in the general time scale of an annual plant generation. As a result, microbial responses to stress may commonly lead to strong and persistent effects on plant natural selection. The role microbial communities play in plant evolution may contribute to variation in selection over space and time as plant populations associating with different microbial communities might respond quite differently to environmental stress. Similarly, microbial legacy effects might cryptically contribute to commonly observed temporal variation in selection (Siepielski *et al*., 2009; Kingsolver & Diamond, 2011). For example, when legacy effects oppose the direction of the effects of stress itself (e.g., as we saw with both fecundity and viability selection on plant flowering time with salt microbes), microbial responses to stress would exacerbate temporal variation in selection for the next generation of plants by increasing the difference in the strength of selection or changing the direction of selection during vs. post stress.

While we focused on selection, evolutionary responses are ultimately also affected by the expression of genetic variation and covariation among traits. Variation in microbial community composition affects these properties as well (O’Brien *et al*. 2019; Bolin, 2023). The net effect microbes play in plant evolution will be determined by all three components (selection, genetic variation, and genetic covariances).

### Conclusions

Overall, we showed that soil microbial responses to stress can contribute to natural selection on plant traits and the opportunity for selection (*I*), potentially altering the evolutionary trajectory of plant populations. Our findings suggest that (1) soil microbial community responses to stress may generally make stress a weaker selective agent than we might expect because they commonly counteract the effects of stress itself, and (2) the evolutionary effects of stress might differ widely depending on the microbial communities present and the responsiveness of those communities to stress. In general, soil microbes and their dynamic responses to environmental change may meaningfully contribute not only to plant ecology, but also to plant evolution.

## ACKNOWLEDGMENTS

LGB was supported as an NSF Graduate Research Fellow. We thank Jeffrey Conner and José Waterton for statistical advice; Maddie Gellinger, Evan Lacey, Ashley Kovach-Hammons, and Trevor Gress for help with data collection, application of stress treatments, and harvest; John Lemon and Tom Pirtle for glasshouse assistance; Susan Magnoli for seeds; Regina Baucom, Mia Howard, Leonie Moyle, Heather Reynolds, Jay Lennon, the Lau and Baucom labs, and two anonymous reviewers for thoughtful comments on this manuscript; and Sarah Fitzpatrick, Jeffrey Conner, and the Fitzpatrick, Conner, and Lau labs for designing and implementing the field experiment from which we sourced soil microbial inocula. This is KBS contribution #2359.

## SUPPORTING INFORMATION

The following Supporting Information is available for this article:

**Methods S1** Statistical methods testing whether effects of stress and microbe history were mediated by changes to the competitive environment

**Table S1** Factors influencing mesocosm productivity

**Table S2** Factors influencing plant linear viability and fecundity selection differentials for specific leaf area (SLA)

**Table S3** Factors influencing plant linear viability and fecundity selection differentials for flowering time

**Table S4** Factors influencing plant linear viability and fecundity selection gradients

**Table S5** Factors influencing plant quadratic viability and fecundity selection differentials for specific leaf area (SLA)

**Table S6** Factors influencing plant quadratic viability and fecundity selection differentials for flowering time

**Table S7** Factors influencing plant quadratic viability and fecundity selection gradients

**Table S8** Factors influencing plant opportunity for selection (*I*), mean absolute fitness ([inline]), and variance in absolute fitness (*σ^2^_W_*)

**Fig. S1** The contemporary environment and microbe history influenced total mesocosm productivity

**Fig. S2** Effects of stress, microbial responses to stress, and microbial legacy effects of stress on plant selection gradients were similar to effects for selection differentials

**Fig. S3** Plots showing fecundity selection for plant specific leaf area (SLA)

**Fig. S4** Plots showing viability selection for plant specific leaf area (SLA)

**Fig. S5** Plots showing viability selection for plant flowering time

**Fig. S6** Plots showing fecundity selection for plant flowering time

## Notes

### Competing Interest Statement

The authors have declared no competing interest.

