## Supplement for "Microbial responses to stress cryptically alter natural selection on plants"

The following Supporting Information is available for this article:

**Methods S1** Statistical methods testing whether effects of stress and microbe history were mediated by changes to the competitive environment

**Table S1** Factors influencing mesocosm productivity

**Table S2** Factors influencing plant linear viability and fecundity selection differentials for specific leaf area (SLA)

**Table S3** Factors influencing plant linear viability and fecundity selection differentials for flowering time

**Table S4** Factors influencing plant linear viability and fecundity selection gradients

**Table S5** Factors influencing plant quadratic viability and fecundity selection differentials for specific leaf area (SLA)

**Table S6** Factors influencing plant quadratic viability and fecundity selection differentials for flowering time

**Table S7** Factors influencing plant quadratic viability and fecundity selection gradients

**Table S8** Factors influencing plant opportunity for selection (*I*), mean absolute fitness ($\underline{W}$*^2^*), and variance in absolute fitness (*σ^2^_W_*)

**Fig. S1** The contemporary environment and microbe history influenced total mesocosm productivity

**Fig. S2** Effects of stress, microbial responses to stress, and microbial legacy of stress on plant selection gradients were similar to effects for selection differentials

**Fig. S3** Plots showing fecundity selection for plant specific leaf area (SLA)

**Fig. S4** Plots showing viability selection for plant specific leaf area (SLA)

**Fig. S5** Plots showing viability selection for plant flowering time

**Fig. S6** Plots showing fecundity selection for plant flowering time

**Methods S1** Statistical methods testing whether effects of stress and microbe history were mediated by changes to the competitive environment

In addition to directly altering natural selection, stress and microbe history may affect selection by altering the competitive environment (e.g., by increasing mortality). For example, all three contemporary stressors reduced mesocosm productivity (i.e., the total biomass produced in each mesocosm), with salt stress reducing productivity the most, followed by herbicide, then herbivory stress (contemp. env.: *F*_3,80_ = 22.5, *P* < 0.001; Table S1; Fig. S1). Microbes also influenced mesocosm productivity, such that herbicide microbes (i.e., microbes from the field herbicide treatment) reduced mesocosm productivity relative to salt microbes (microbe history: *F*_3,80_ = 3.2, *P* = 0.03; Table S1; Fig. S1). The reduced competition in these treatments could then alter natural selection on plant traits. Therefore, in addition to the fecundity selection models described above that estimate the total (both direct and competition-mediated) effects of stress and microbes on selection, we fit additional models that estimate the direct effects alone by controlling for variation in plant productivity across mesocosms. To do this, we first fit a linear mixed model that included aboveground biomass as the response variable, mesocosm productivity as the predictor, and mesocosm as a random effect. We then added each plant’s residual value from this model to the global mean aboveground biomass value to return biomass to its original scale. These “detrended” biomass data represent plant biomass after removing the effect of competition on biomass. We did not ln-transform the detrended data because some detrended biomass values were negative, for which the natural logarithm would be undefined. Finally, we relativized the detrended dataset globally for selection coefficient estimation. To test for the effects of stress and microbes independent of effects caused by changes in the competitive environment, we then fit identical models as described above except we included this detrended biomass value as the response variable. Controlling for mesocosm productivity did not cause any significant terms to lose significance, suggesting that the effects of stress and microbe history on plant natural selection were not mediated by changes in the competitive environment (Tables S2-S7).

**Table S1** Results of a linear model testing the effects of contemporary environment (“Contemp. Env.”) and microbe history (“Microbes”) on mesocosm productivity (total pot aboveground biomass).

| **Term** | **df** | ***F*** | ***P*** |
| --- | --- | --- | --- |
| Contemp. Env. | 3 | 22.515 | **< 0.001** |
| Microbes | 3 | 3.157 | **0.029** |
| Microbes × Contemp. Env. | 9 | 1.142 | 0.344 |
| Residuals | 80 |  |  |

**Table S2** Results of a linear model testing the effects of contemporary environment (“Contemp. Env.”) and microbe history (“Microbes”) on plant linear viability and fecundity selection differentials for specific leaf area (SLA).

|  | Viability | | | Fecundity | | | Fecundity  (after controlling for plant productivity) | | |
| --- | --- | --- | --- | --- | --- | --- | --- | --- | --- |
| **Term** | **df** | ***𝝌^2^*** | ***P*** | **df** | ***F*** | ***P*** | **df** | ***F*** | ***P*** |
| Microbes | 1 | 1.345 | 0.246 | 1 | 0.674 | 0.413 | 1 | < 0.001 | 0.994 |
| Contemp. Env. | 1 | 66.620 | **< 0.001** | 1 | 19.686 | **< 0.001** | 1 | 6.837 | **0.01** |
| SLA | 1 | 0.350 | 0.554 | 1 | 34.919 | **< 0.001** | 1 | 31.442 | **< 0.001** |
| Microbes × Contemp. Env. | 1 | 0.237 | 0.626 | 1 | 2.686 | 0.103 | 1 | 0.104 | 0.747 |
| Contemp. Env. × SLA | 1 | 4.394 | **0.036** | 1 | 5.282 | **0.023** | 1 | 17.652 | **< 0.001** |
| Microbes × SLA | 1 | 6.169 | **0.013** | 1 | 0.024 | 0.877 | 1 | 0.112 | 0.738 |
| Microbes **×** Contemp. Env. **×** SLA | 1 | 0.362 | 0.547 | 1 | 0.099 | 0.753 | 1 | 0.004 | 0.952 |
| Residuals |  |  |  | 172 |  |  | 172 |  |  |

Notes: Probability of survival to flower (proportion of surviving individuals in each family) was the response variable for viability selection, ln-transformed aboveground biomass was the response variable for fecundity selection, and aboveground biomass detrended by mesocosm productivity (see Methods S1) was the response variable for fecundity selection after controlling for plant productivity. For viability selection, each data point was weighted by the number of individuals in each family × treatment combination (range: 1-3 due to early seedling mortality; mode: 3). We used family mean trait values, and standardized plant trait values (SLA) to a mean of 0 and a standard deviation of 1. Significant p-values (*P* < 0.05) are bolded.

**Table S3** Results of a generalized linear model testing the effects of contemporary environment (“Contemp. Env.”) and microbe history (“Microbes”) on plant linear viability and fecundity selection differentials for flowering time.

|  | Viability | | | Fecundity | | | Fecundity  (after controlling for plant productivity) | | |
| --- | --- | --- | --- | --- | --- | --- | --- | --- | --- |
| **Term** | **df** | ***𝝌^2^*** | ***P*** | **df** | ***F*** | ***P*** | **df** | ***F*** | ***P*** |
| Microbes | 3 | 15.058 | **0.002** | 3 | 3.923 | **0.009** | 3 | 5.338 | **0.001** |
| Contemp. Env. | 3 | 125.852 | **< 0.001** | 3 | 18.652 | **< 0.001** | 3 | 6.283 | **< 0.001** |
| Flowering Time | 1 | 0.035 | 0.852 | 1 | 74.680 | **< 0.001** | 1 | 69.751 | **< 0.001** |
| Microbes × Contemp. Env. | 9 | 34.144 | **< 0.001** | 9 | 1.332 | 0.217 | 9 | 0.717 | 0.693 |
| Contemp. Env. × Flowering Time | 3 | 3.921 | 0.270 | 3 | 2.393 | **0.067** | 3 | 2.215 | **0.085** |
| Microbes × Flowering Time | 3 | 8.233 | **0.041** | 3 | 0.949 | 0.416 | 3 | 1.515 | 0.209 |
| Microbes × Contemp. Env. × Flowering Time | 9 | 9.725 | 0.373 | 9 | 0.724 | 0.687 | 9 | 1.170 | 0.311 |
| Residuals |  |  |  | 650 |  |  | 650 |  |  |

Notes: Probability of survival to flower (proportion of surviving individuals in each family) was the response variable for viability selection, ln-transformed aboveground biomass was the response variable for fecundity selection, and aboveground biomass detrended by mesocosm productivity (see Methods S1) was the response variable for fecundity selection after controlling for plant productivity. For viability selection, each data point was weighted by the number of individuals in each family × treatment combination (range: 1-3 due to early seedling mortality; mode: 3). We used family mean trait values, and standardized plant trait values (SLA) to a mean of 0 and a standard deviation of 1. Significant and marginally significant p-values (*P* < 0.1) are bolded.

**Table S4** Results of a linear model testing the effects of contemporary environment (“Contemp. Env.”) and microbe history (“Microbes”) on plant linear viability and fecundity selection gradients.

|  | Viability | | | Fecundity | | | Fecundity  (after controlling for plant productivity) | | |
| --- | --- | --- | --- | --- | --- | --- | --- | --- | --- |
| **Term** | **df** | ***𝝌^2^*** | ***P*** | **df** | ***F*** | ***P*** | **df** | ***F*** | ***P*** |
| Microbes | 1 | 1.419 | 0.234 | 1 | 2.310 | 0.130 | 1 | 0.152 | 0.697 |
| Contemp. Env. | 1 | 69.045 | **< 0.001** | 1 | 14.911 | **< 0.001** | 1 | 11.077 | **0.001** |
| Flowering Time | 1 | 1.858 | 0.173 | 1 | 6.789 | **0.010** | 1 | 6.308 | **0.013** |
| SLA | 1 | 3.742 | **0.053** | 1 | 34.708 | **< 0.001** | 1 | 28.685 | **< 0.001** |
| Microbes × Contemp. Env. | 1 | 0.032 | 0.858 | 1 | 4.932 | **0.028** | 1 | 0.015 | 0.902 |
| Contemp. Env. × Flowering Time | 1 | 0.025 | 0.873 | 1 | 1.346 | 0.248 | 1 | 3.082 | **0.081** |
| Contemp. Env. × SLA | 1 | 4.731 | **0.030** | 1 | 2.792 | **0.097** | 1 | 11.447 | **< 0.001** |
| Microbes × Flowering Time | 1 | 2.319 | 0.128 | 1 | 0.094 | 0.760 | 1 | 0.633 | 0.427 |
| Microbes × SLA | 1 | 0.778 | 0.378 | 1 | 0.285 | 0.594 | 1 | 0.184 | 0.669 |
| Microbes × Contemp. Env. × Flowering Time | 1 | 0.988 | 0.320 | 1 | 0.367 | 0.545 | 1 | 0.792 | 0.375 |
| Microbes × Contemp. Env. × SLA | 1 | 0.155 | 0.694 | 1 | 0.028 | 0.868 | 1 | 0.260 | 0.611 |
| Residuals |  |  |  | 161 |  |  | 161 |  |  |

Notes: Probability of survival to flower (proportion of surviving individuals in each family) was the response variable for viability selection, ln-transformed aboveground biomass was the response variable for fecundity selection, and aboveground biomass detrended by mesocosm productivity (see Methods S1) was the response variable for fecundity selection after controlling for plant productivity. For viability selection, each data point was weighted by the number of individuals in each family × treatment combination (range: 1-3 due to early seedling mortality; mode: 3). We used family mean trait values, and standardized plant trait values (SLA) to a mean of 0 and a standard deviation of 1. Significant and marginally significant p-values (*P* < 0.1) are bolded.

**Table S5** Results of a generalized linear model testing the effects of contemporary environment (“Contemp. Env.”) and microbe history (“Microbes”) on plant quadratic viability and fecundity selection differentials for specific leaf area (SLA).

|  | Viability | | | Fecundity | | | Fecundity  (after controlling for plant productivity) | | |
| --- | --- | --- | --- | --- | --- | --- | --- | --- | --- |
| **Term** | **df** | ***𝝌^2^*** | ***P*** | **df** | ***F*** | ***P*** | **df** | ***F*** | ***P*** |
| Microbes | 1 | 1.138 | 0.286 | 1 | 0.741 | 0.390 | 1 | < 0.001 | 0.988 |
| Contemp. Env. | 1 | 70.300 | **< 0.001** | 1 | 21.935 | **< 0.001** | 1 | 5.487 | **0.020** |
| SLA | 1 | 7.075 | **0.008** | 1 | 30.160 | **< 0.001** | 1 | 26.106 | **< 0.001** |
| SLA^2^ | 1 | 10.589 | **0.001** | 1 | 2.427 | 0.121 | 1 | 1.754 | 0.187 |
| Microbes × Contemp. Env. | 1 | 0.440 | 0.507 | 1 | 1.513 | 0.220 | 1 | 0.264 | 0.608 |
| Contemp. Env. × SLA | 1 | 9.767 | **0.002** | 1 | 1.790 | 0.183 | 1 | 7.064 | **0.009** |
| Microbes × SLA | 1 | 2.591 | 0.107 | 1 | 0.009 | 0.924 | 1 | 0.055 | 0.815 |
| Contemp. Env. × SLA^2^ | 1 | 0.191 | 0.662 | 1 | 1.155 | 0.284 | 1 | 0.101 | 0.752 |
| Microbes **×** SLA^2^ | 1 | 0.183 | 0.669 | 1 | 1.481 | 0.225 | 1 | 0.191 | 0.662 |
| Microbes **×** Contemp. Env. **×** SLA | 1 | 1.812 | 0.178 | 1 | 0.018 | 0.894 | 1 | 0.049 | 0.824 |
| Microbes **×** Contemp. Env. **×** SLA^2^ | 1 | 2.350 | 0.125 | 1 | 0.228 | 0.634 | 1 | 0.257 | 0.613 |
| Residuals |  |  |  | 168 |  |  | 168 |  |  |

Notes: Probability of survival to flower (proportion of surviving individuals in each family) was the response variable for viability selection, ln-transformed aboveground biomass was the response variable for fecundity selection, and aboveground biomass detrended by mesocosm productivity (see Methods S1) was the response variable for fecundity selection after controlling for plant productivity. For viability selection, each data point was weighted by the number of individuals in each family × treatment combination (range: 1-3 due to early seedling mortality; mode: 3). We used family mean trait values, and standardized plant trait values (SLA) to a mean of 0 and a standard deviation of 1. Significant and marginally significant p-values (*P* < 0.1) are bolded.

**Table S6** Results of a generalized linear model testing the effects of contemporary environment (“Contemp. Env.”) and microbe history (“Microbes”) on plant quadratic viability and fecundity selection differentials for flowering time.

|  | Viability | | | Fecundity | | | Fecundity  (after controlling for plant productivity) | | |
| --- | --- | --- | --- | --- | --- | --- | --- | --- | --- |
| **Term** | **df** | ***𝝌^2^*** | ***P*** | **df** | ***F*** | ***P*** | **df** | ***F*** | ***P*** |
| Microbes | 3 | 14.170 | **0.003** | 3 | 3.788 | **0.010** | 3 | 5.540 | **< 0.001** |
| Contemp. Env. | 3 | 117.972 | **< 0.001** | 3 | 19.287 | **< 0.001** | 3 | 6.711 | **< 0.001** |
| Flowering Time | 1 | 0.932 | 0.334 | 1 | 43.217 | **< 0.001** | 1 | 48.242 | **< 0.001** |
| Flowering Time^2^ | 1 | 3.216 | **0.073** | 1 | 11.853 | **<0.001** | 1 | 4.417 | **0.036** |
| Microbes × Contemp. Env. | 9 | 35.568 | **< 0.001** | 9 | 1.174 | 0.309 | 9 | 0.769 | 0.645 |
| Contemp. Env. × Flowering Time | 3 | 3.392 | 0.335 | 3 | 2.899 | **0.034** | 3 | 2.838 | **0.037** |
| Microbes × Flowering Time | 3 | 9.995 | **0.019** | 3 | 1.512 | 0.210 | 3 | 2.190 | **0.088** |
| Contemp. Env. × Flowering Time^2^ | 3 | 2.453 | 0.484 | 3 | 1.357 | 0.255 | 3 | 0.883 | 0.450 |
| Microbes × Flowering Time^2^ | 3 | 1.118 | 0.773 | 3 | 0.140 | 0.936 | 3 | 1.110 | 0.345 |
| Microbes × Contemp. Env. × Flowering Time | 9 | 11.210 | 0.262 | 9 | 1.267 | 0.252 | 9 | 2.009 | **0.036** |
| Microbes × Contemp. Env. × Flowering Time^2^ | 9 | 12.231 | 0.196 | 9 | 1.203 | 0.290 | 9 | 1.333 | 0.216 |
| Residuals |  |  |  | 634 |  |  | 634 |  |  |

Notes: Probability of survival to flower (proportion of surviving individuals in each family) was the response variable for viability selection, ln-transformed aboveground biomass was the response variable for fecundity selection, and aboveground biomass detrended by mesocosm productivity (see Methods S1) was the response variable for fecundity selection after controlling for plant productivity. For viability selection, each data point was weighted by the number of individuals in each family × treatment combination (range: 1-3 due to early seedling mortality; mode: 3). We used family mean trait values, and standardized plant trait values (flowering time) to a mean of 0 and a standard deviation of 1. Significant and marginally significant p-values (*P* < 0.1) are bolded.

**Table S7** Results of a generalized linear model testing the effects of contemporary environment (“Contemp. Env.”) and microbe history (“Microbes”) on plant quadratic viability and fecundity selection gradients.

|  | Viability | | | Fecundity | | | Fecundity  (after controlling for plant productivity) | | |
| --- | --- | --- | --- | --- | --- | --- | --- | --- | --- |
| **Term** | **df** | ***𝝌^2^*** | ***P*** | **df** | ***F*** | ***P*** | **df** | ***F*** | ***P*** |
| Microbes | 1 | 0.851 | 0.356 | 1 | 2.813 | **0.096** | 1 | 0.350 | 0.555 |
| Contemp. Env. | 1 | 70.479 | **< 0.001** | 1 | 16.721 | **< 0.001** | 1 | 8.923 | **0.003** |
| Flowering Time | 1 | 2.952 | **0.086** | 1 | 6.158 | **0.014** | 1 | 5.679 | **0.018** |
| SLA | 1 | 4.440 | **0.035** | 1 | 33.264 | **< 0.001** | 1 | 24.601 | **< 0.001** |
| Flowering Time^2^ | 1 | 1.950 | 0.163 | 1 | 0.116 | 0.734 | 1 | 0.874 | 0.351 |
| SLA^2^ | 1 | 4.089 | **0.043** | 1 | 2.335 | 0.129 | 1 | 1.381 | 0.242 |
| Microbes × Contemp. Env. | 1 | 0.035 | 0.851 | 1 | 2.724 | 0.101 | 1 | 0.162 | 0.688 |
| Contemp. Env. × Flowering Time | 1 | 0.073 | 0.788 | 1 | 1.472 | 0.227 | 1 | 4.439 | **0.037** |
| Contemp. Env. × SLA | 1 | 9.026 | **0.003** | 1 | 0.621 | 0.432 | 1 | 3.971 | **0.048** |
| Microbes × Flowering Time | 1 | 2.312 | 0.128 | 1 | 0.029 | 0.866 | 1 | 0.005 | 0.944 |
| Microbes × SLA | 1 | 2.321 | 0.128 | 1 | 0.105 | 0.746 | 1 | 0.160 | 0.690 |
| Contemp. Env. × Flowering Time^2^ | 1 | 2.491 | 0.114 | 1 | 0.328 | 0.567 | 1 | 0.268 | 0.605 |
| Contemp. Env. × SLA^2^ | 1 | 0.651 | 0.420 | 1 | 1.918 | 0.168 | 1 | 0.324 | 0.570 |
| Microbes × Flowering Time^2^ | 1 | 0.049 | 0.825 | 1 | 0.013 | 0.911 | 1 | 0.003 | 0.958 |
| Microbes × SLA^2^ | 1 | 0.256 | 0.613 | 1 | 0.808 | 0.370 | 1 | 0.034 | 0.855 |
| Microbes × Contemp. Env. × Flowering Time | 1 | 0.433 | 0.510 | 1 | 0.528 | 0.469 | 1 | 0.557 | 0.457 |
| Microbes × Contemp. Env. × SLA | 1 | 0.757 | 0.384 | 1 | 0.006 | 0.938 | 1 | 0.008 | 0.927 |
| Microbes × Contemp. Env. × Flowering Time^2^ | 1 | 0.008 | 0.929 | 1 | 0.220 | 0.640 | 1 | 0.043 | 0.836 |
| Microbes **×** Contemp. Env. **×** SLA^2^ | 1 | 2.030 | 0.154 | 1 | 0.363 | 0.548 | 1 | 0.649 | 0.422 |
| Residuals |  |  |  | 153 |  |  | 153 |  |  |

Notes: Probability of survival to flower (proportion of surviving individuals in each family) was the response variable for viability selection, ln-transformed aboveground biomass was the response variable for fecundity selection, and aboveground biomass detrended by mesocosm productivity (see Methods S1) was the response variable for fecundity selection after controlling for plant productivity. For viability selection, each data point was weighted by the number of individuals in each family × treatment combination (range: 1-3 due to early seedling mortality; mode: 3). We used family mean trait values, and standardized plant trait values (flowering time and SLA) to a mean of 0 and a standard deviation of 1. Significant and marginally significant p-values (*P* < 0.1) are bolded.

**Table S8** Results of a linear model testing the effects of contemporary environment (“Contemp. Env.”) and microbe history (“Microbes”) on plant opportunity for selection (*I*), mean absolute fitness squared

($\underline{W}$*^2^*), and variance in absolute fitness (*σ^2^_W_*).

|  | *I* | | | $\underline{W}$*^2^* | | | *σ^2^_W_* | | |
| --- | --- | --- | --- | --- | --- | --- | --- | --- | --- |
| **Term** | **df** | ***F*** | ***P*** | **df** | ***F*** | ***P*** | **df** | ***F*** | ***P*** |
| Microbes | 3 | 6.43 | **< 0.001** | 3 | 1.58 | 0.201 | 3 | 0.57 | 0.635 |
| Contemp. Env. | 3 | 5.16 | **0.003** | 3 | 22.39 | **< 0.001** | 3 | 6.25 | **< 0.001** |
| Microbes **×** Contemp. Env. | 9 | 1.10 | 0.376 | 9 | 1.09 | 0.377 | 9 | 0.29 | 0.976 |
| Residuals | 80 |  |  | 80 |  |  | 80 |  |  |

Notes: Variance in relative fitness was the response variable for *I* model.

**Fig. S1** The contemporary environment and microbe history influenced total mesocosm productivity.

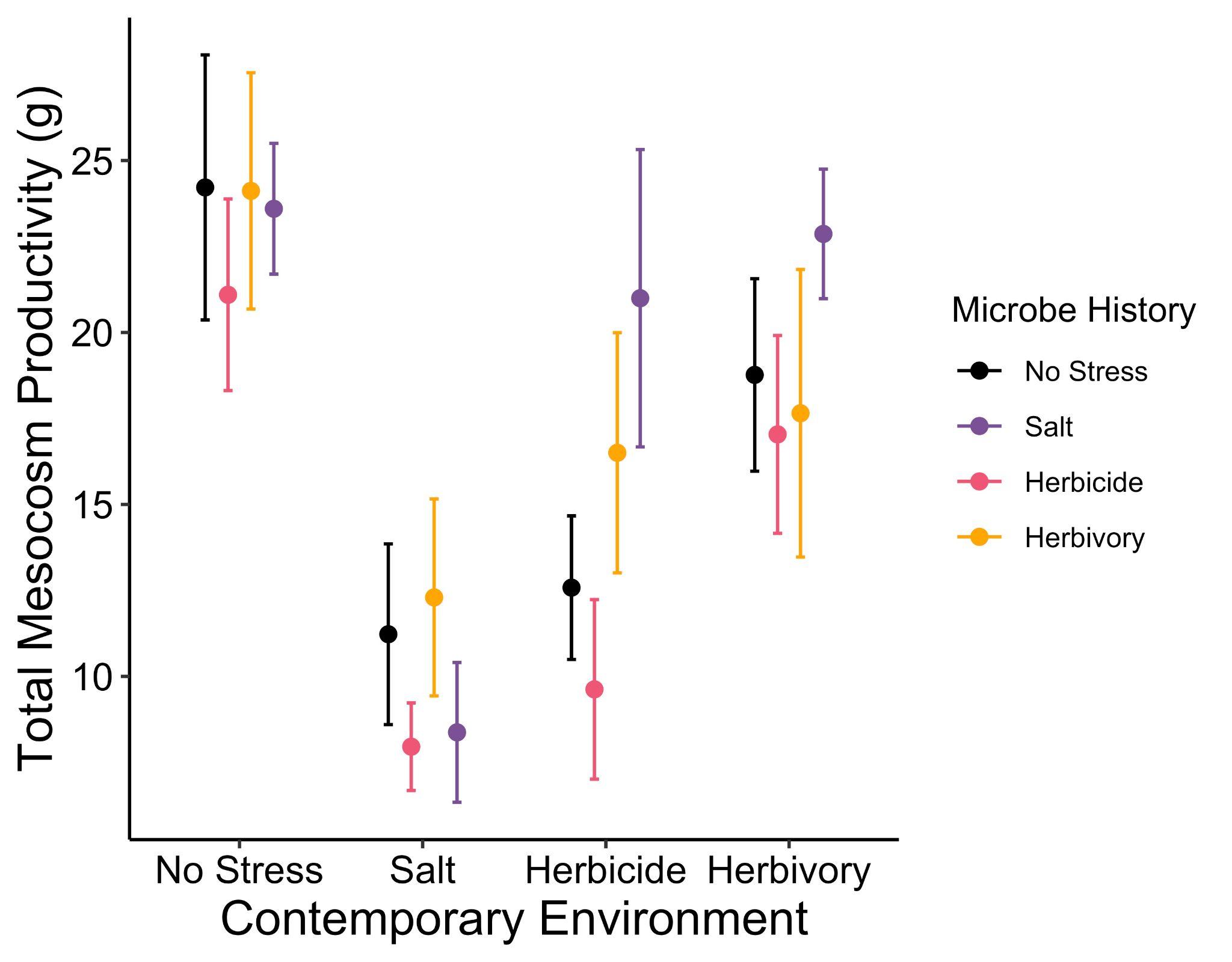

**Figure S2** Effects of stress, microbial responses to stress, and microbial legacy of stress on plant selection gradients were similar to effects for selection differentials (Fig. 4 in the main text; we were limited to salt stress for selection gradients as described in Methods). Selection on plant (a) SLA and (b) flowering time can be affected by the stress environment itself (black bars), by microbial community responses to stress (gray bars), or by microbial legacy effects of past stress (white bars). Legacy effects occur when stress selects for microbial communities that alter selection even after the stress has ceased (i.e., in non stressful contemporary environments). Effects of stress, microbial response to stress, and legacy of microbial response to stress were calculated as described in equations 1, 2, and 3. Values above zero indicate that an effect caused selection differentials to become more positive (or less negative), and values below zero indicate an effect caused selection to become more negative (or less positive), as depicted in the cartoon graphs on the y-axis in (a). Fecundity and viability selection are shown on the left and right side of each panel, respectively, and contemporary stress environments are shown across the x-axis.

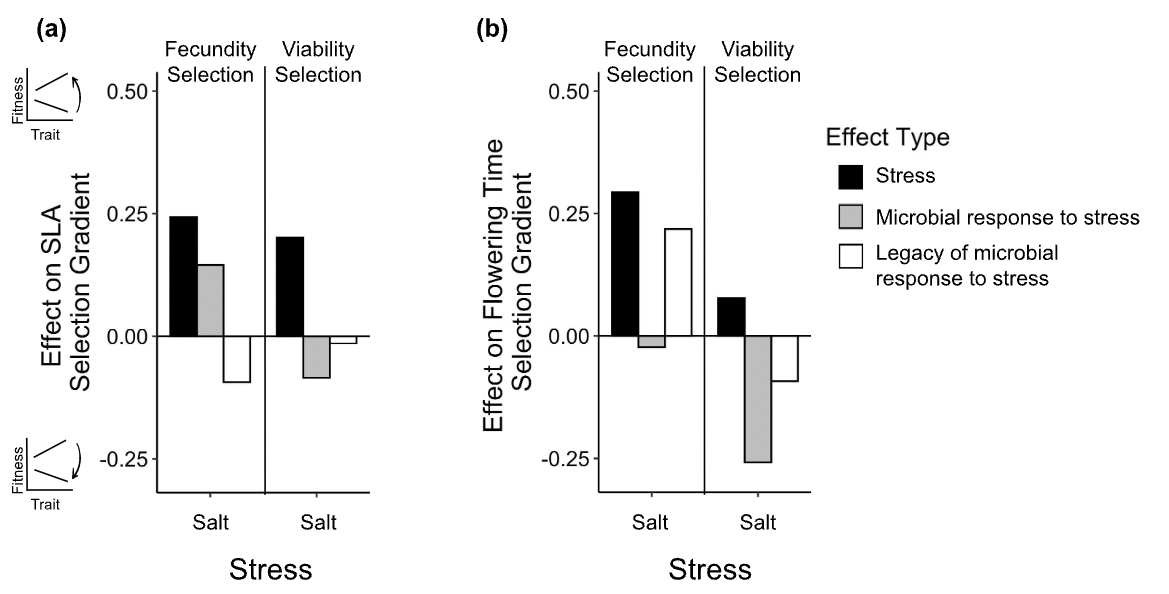

**Figure S3** Plots showing fecundity selection for plant specific leaf area (SLA) in (a) non-stressful environments and (b) salt stress. Aboveground biomass was relativized by mean aboveground biomass, and each point represents the family mean. Standardized SLA was standardized by the standard deviation globally (across all treatments), and represents the family mean trait value. “No stress” and “salt” microbe history treatments represent soil microbes from field plots where plants were unstressed (black) or salt-stressed (purple), respectively.

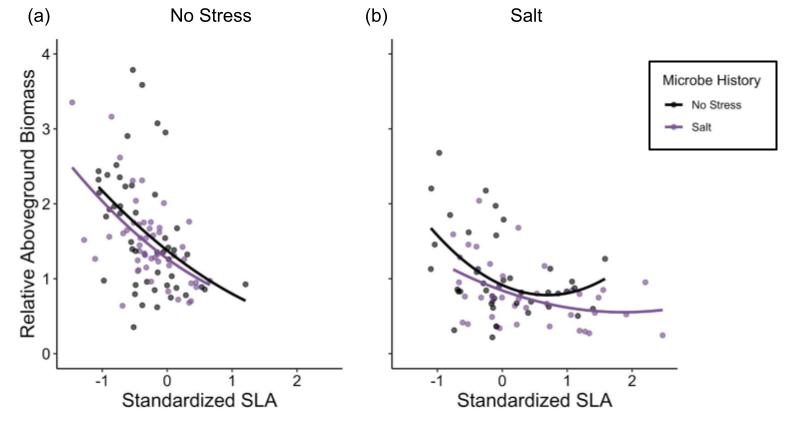

**Figure S4** Plots showing viability selection for plant specific leaf area (SLA) in (a) non stressful environments and (b) salt stress. Survival probability is the proportion of individuals in a maternal family that survived under each contemporary environment × microbe history treatment. Points are jittered vertically for clarity. Standardized SLA was standardized by the standard deviation globally (across all treatments), and represents the family mean trait value. “No stress” and “salt” microbe history treatments represent soil microbes from field plots where plants were unstressed (black) or salt-stressed (purple), respectively.

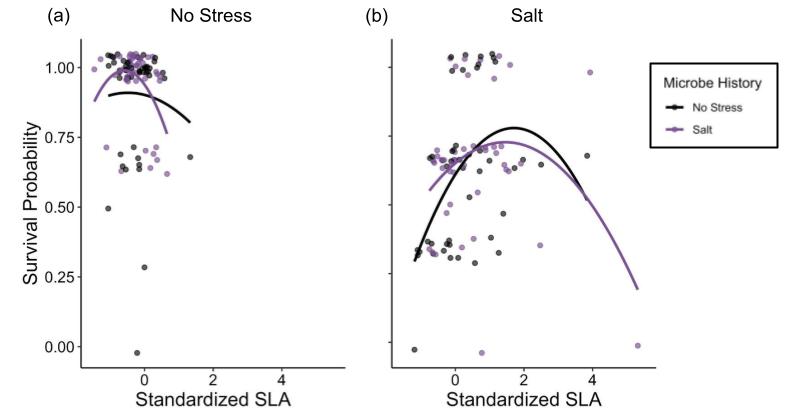

**Figure S5** Plots showing viability selection for plant flowering time in (a) non stressful environments, (b) salt stress, (c) herbicide stress, and (d) herbivory stress. Survival probability is the proportion of individuals in a maternal family that survived under each contemporary environment × microbe history treatment. Points are jittered vertically for clarity. Standardized flowering time was standardized by the standard deviation globally (across all treatments), and represents the family mean trait value. “No stress,” “salt”, “herbicide,” and “herbivory” microbe history treatments represent soil microbes from field plots where plants were unstressed (black), salt-stressed (purple), herbicide-stressed (pink), and herbivory-stressed (yellow), respectively.

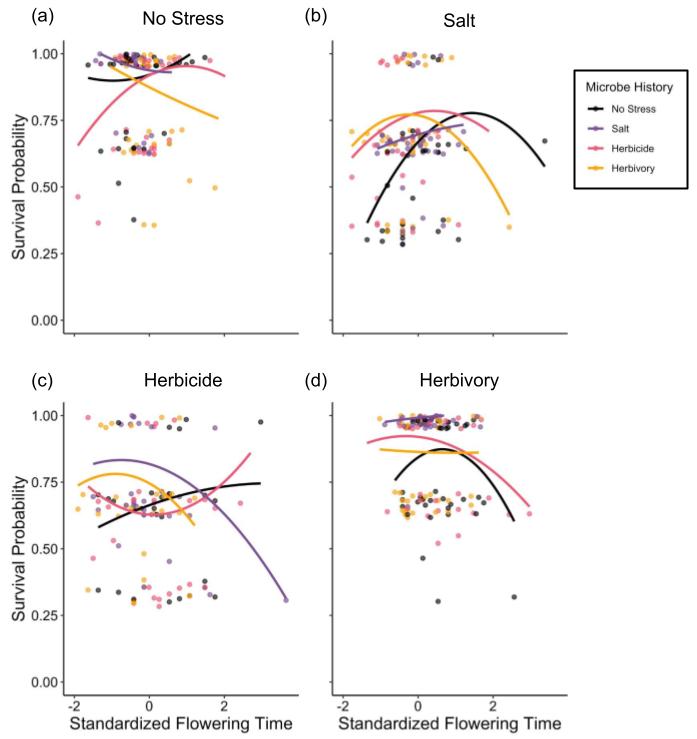

**Figure S6** Plots showing fecundity selection for plant flowering time in (a) non stressful environments, (b) salt stress, (c) herbicide stress, and (d) herbivory stress. Aboveground biomass was relativized by mean aboveground biomass, and each point represents the family mean. Standardized flowering time was standardized by the standard deviation globally (across all treatments), and represents the family mean trait value. “No stress,” “salt”, “herbicide,” and “herbivory” microbe history treatments represent soil microbes from field plots where plants were unstressed (black), salt-stressed (purple), herbicide-stressed (pink), and herbivory-stressed (yellow), respectively.

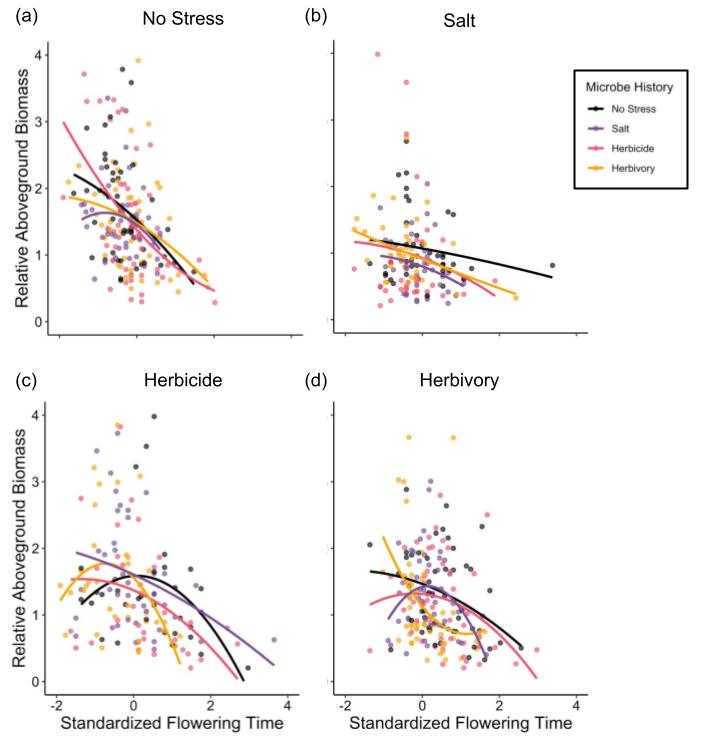
